# Experimental evolution of collective β-lactam resistance in *Escherichia coli* via activation of a dormant outermembrane porin

**DOI:** 10.64898/2026.08.19.745717

**Authors:** Abraham Ochoa-Guerrero, Joost Hollander, Rotem Gross, Aditi Batra, Josia Pool, Francisca C. Reyes Márquez, Arjen C. van de Peppel, Joachim Krug, J. Arjan G.M. de Visser

## Abstract

Understanding the mechanisms that drive antibiotic resistance is relevant for both evolutionary theory and the design of effective drug therapies. A specific challenge are collective resistance mechanisms, where bacterial populations survive drug concentrations that kill individual bacteria. Here, we explore the evolvability of collective resistance mechanisms in bacterial strains expressing antibiotic-degrading β-lactamases with different levels of privatization. Four strains of *Escherichia coli*, with or without outermembrane porin OmpF to affect drug permeability and expressing either a low or high-activity periplasmic β-lactamase, were subjected to lineage selection in a gradient of the drug cefotaxime. Strains with a low-activity enzyme increased cell-level resistance, while strains with low permeability, and hence a more private β-lactamase function, increased collective resistance. Remarkably, increased collective resistance in the strain with a private high-activity β-lactamase came with decreased cell-level resistance. This tradeoff was primarily caused by the activation of a dormant outermembrane porin, NmpC, via the excision of an insertion sequence. Increased drug permeability through NmpC explained both its lower cell-level resistance and its greater collective resistance through faster drug removal and growth recovery via enhanced filamentation at high cell density. The recovery advantage of the NmpC mutant also explained its initial invasion within the ancestral population, suggesting that drug permeability is a readily evolvable collective-resistance mechanism in bacteria with high-activity β-lactamases. Our findings highlight the role of filamentation and drug permeability in β-lactamase-mediated collective resistance to these widely used drugs.

## Introduction

The rapid development of antibiotic resistance in pathogenic bacteria is a global public health concern, contributing to an increasing number of deaths each year.^1,2^ Particularly, bacterial resistance to β-lactam antibiotics poses a serious threat due to the clinical importance of these drugs.^3^

β-Lactam antibiotics target penicillin-binding proteins (PBPs) located in the periplasmic space of gram-negative bacteria. PBPs belong to an enzyme family involved in the biogenesis of the bacterial envelope. Binding of β-lactam antibiotics to PBPs disrupt bacterial cell wall synthesis preventing bacterial proliferation.^4^ Among the different PBPs expressed by bacteria, β-lactams, particularly cephalosporins, primarily bind to PBP3.^5^ This enzyme plays a central role in the divisome, a multienzyme complex responsible for bacterial septation.^6,7^ Consequently, inhibition of PBP3 activity induces filamentous growth, a phenomenon that has been suggested to enhance bacterial survival in toxic environments.^8–10^

Bacteria have devolved various resistance mechanisms to β-lactams.^11,12^ Resistance to high antibiotic concentrations is often associated with the production of periplasmic β-lactamase enzymes that inactivate β-lactams by hydrolyzing their β-lactam ring.^13,14^ In addition to enzymatic degradation, other common resistance mechanisms identified in *E. coli* include the reduction of periplasmic antibiotic concentration via the upregulation of efflux pumps ^15,16^ or the reduction of antibiotic uptake through the repression of outer membrane porins.^17,18^

Importantly, the evolution of resistance mechanisms is driven not only by their effects on individual bacterial survival but also by collective benefits. For example, β-lactamase-mediated drug degradation increases with cell density, enabling bacterial populations to survive β-lactam concentrations that are lethal at low cell density.^19,20^ Such collective resistance can be observed as an “inoculum effect”, where larger bacterial populations require higher minimal inhibitory concentrations (MIC) of the drug.^21–25^ For example, enzymatic degradation of antibiotics in the periplasm helps detoxify the extracellular environment, allowing even antibiotic-sensitive cells within the population to survive.^26–32^

Because β-lactamases reside in the periplasm, intracellular drug degradation depends on the influx of these molecules across the outermembrane.^33^ β-Lactam antibiotics penetrate the outer membrane of gram-negative bacteria primarily through porin-mediated diffusion. This process is relatively slow and often constitutes the rate-limiting step in drug access in the periplasm.^34,35^ Hence, the release of β- lactamases as a public-good into the extracellular environment may faster remove the drug,^19,36^ e.g. via outer membrane vesicles.^37^

β-Lactamases are also naturally released upon cell death, creating the possibility of enhanced collective survival through altruistic cell lysis.^38^ Support for this idea comes from the observation that β- lactamase-producing strains challenged with β-lactam antibiotics exhibit non-monotonic growth curves consisting of an initial growth phase, during which bacteria elongate and grow as filaments—most likely due to the inhibition of PBP3—followed by a decay phase that might facilitate β-lactamase release and accelerate antibiotic degradation, and finally a regrowth phase once the medium has been sufficiently detoxified.^39,40^ Studies with engineered strains expressing a cytoplasmic β-lactamase and inducible suicide module, suggest that optimal lysis rates for collective survival exist under conditions where the benefit provided by the public good is sufficiently high and opportunities for invasion of non-producing cheaters are limited.^38,41^ However, the conditions for the *de novo* evolution of such self-destructive cooperation in bacterial communities are still unclear.^41–43^

In this study, we sought to select spontaneous mutants of *Escherichia coli* that exhibit altruistic collective resistance to the cephalosporin cefotaxime (CTX). To this end, we used four *E. coli* variants that differed in their ability to share intracellular β-lactamase activity as a public good. This was achieved by varying their outer membrane permeability to CTX through the presence or absence of the outer membrane porin OmpF, as well as by altering the catalytic efficiency of their β-lactamases against CTX (Figure 1A). To maximize selection for collective resistance, we used a lineage selection protocol in a gradient of CTX concentrations (Figure 1B). Based in previous observations,^39^ we expected the capacity to evolve β-lactamase-mediated altruistic resistance to be the lowest for the permeable strain expressing low-activity β-lactamase and highest for the non-permeable strain expressing the high-activity β-lactamase. Whole-genome sequencing analysis, together with several phenotypic analyses (Figure 1C)—including the inoculum effect on MIC, growth kinetics in the presence of antibiotics, characterization of cell morphology using microscopy and flow cytometry, and measurement of environmental CTX degradation by HPLC—provided strong evidence that initially low permeability to CTX, combined with high catalytic efficiency, drives the evolution of altruistic collective resistance through increased antibiotic permeability.

**Figure 1.**
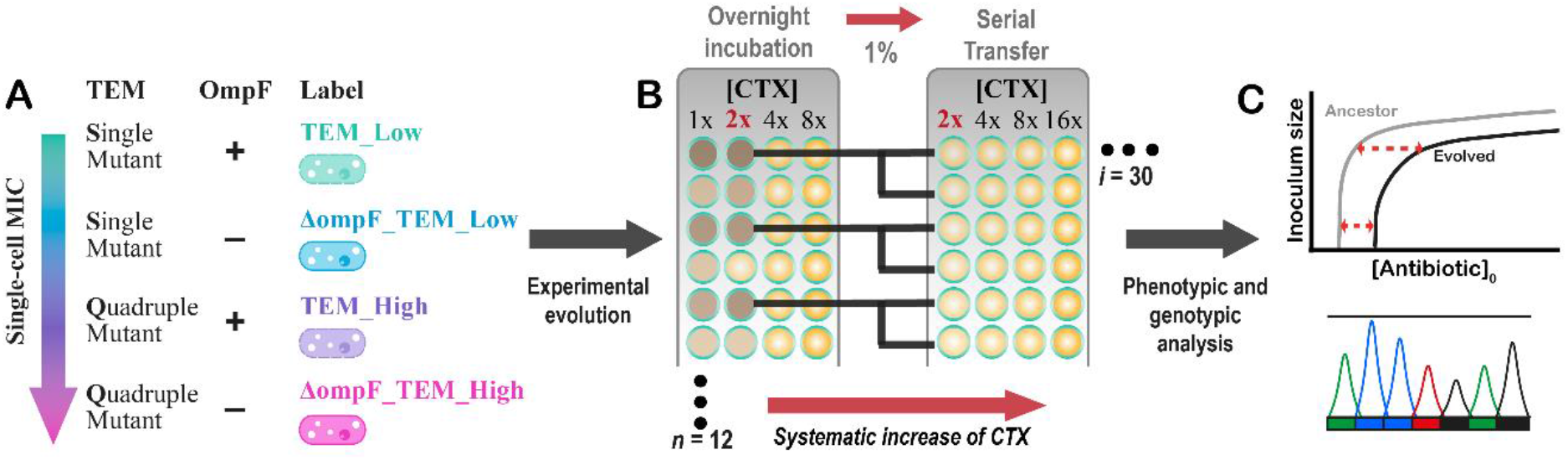
Method overview for the experimental evolution of altruistic collective resistance of four different β- lactamase-producer strains of *E. coli*. (A) The four strains were selected based on CTX permeability and β-lactamase catalytic efficiency, which determine their potential to share intracellular β-lactamase activity. (B) Selection strategy used for each transfer cycle during the experiment of evolution, each strain was subjected independently to the same protocol (see Methods for details). (C) After ∼160 generations, the evolved lineages were characterized phenotypically and genotypically to elucidate the collective mechanisms underlying the increased resistance to CTX treatment.

## Results

### Evolutionary dynamics

We carried out an evolution experiment aimed at exploring the evolvability of collective resistance to CTX treatment in four strains of *E. coli* REL606 with different potential to share their intracellular β- lactamase activity: strains either express outermembrane porin OmpF or not due to its deletion (Δ*ompF*), and express either low-activity TEM-Low (TEM-1 with single mutation: G238S) or high-activity TEM-High (TEM-1 with four mutations: E104K, M182T, G238S and T265M) β-lactamase, (Figure 1A and Supplementary Table S1, SI). We hypothesize that the more efficient the β-lactamase produced and the lower the permeability of the outer membrane to CTX molecules, the more likely a bacterial lineage is to evolve self-destructive mechanisms that release β-lactamases into the extracellular space, thereby indirectly benefiting other members of the population and promoting collective resistance.

To increase the chances of selection for population-level resistance mechanisms, we developed an experimental evolution protocol that combines selection at the individual and population levels. Briefly, we subjected 12 replicate lineages of each strain to a lineage selection protocol for increased MIC at high cell density (∼5 x 10^6^ cells/mL) in a gradient of four two-fold increasing CTX concentrations (48 replicate populations per strain), allowing the six lineages with highest MIC (or highest yield in case of no variation in MIC) to each found two new lineages (Figure 1B; see details in the Methods). For each strain, we tracked the six evolved lineages during 30 transfers (∼160 generations), at the end all final six were derived from a common ancestral lineage, indicating their relatedness (Supplementary Figure S2 and S3). CTX tolerance increased by 128-fold in the low-activity β-lactamase producer lineages (TEM-Low and Δ*ompF*_TEM-Low) compared with a 4-fold increase in the high-activity producer lineages (TEM-High and Δ*ompF*_TEM-High). The low- and high-activity β-lactamase producers reached their maximum tolerated CTX concentrations after seven and nine transfers, respectively (Supplementary Figure S2 and S3).

### Changes in individual and collective resistance

To evaluate the increase in individual and collective CTX resistance in the evolved lineages, we assessed the dependence of the MIC on the initial bacterial population density (inoculum size) based on the Minimal Survival Inoculum curves (MSI curves, Figure 2A and S4), using the model of Geyrhofer et al. (2023) shown in equation 1.

**Figure 2.**
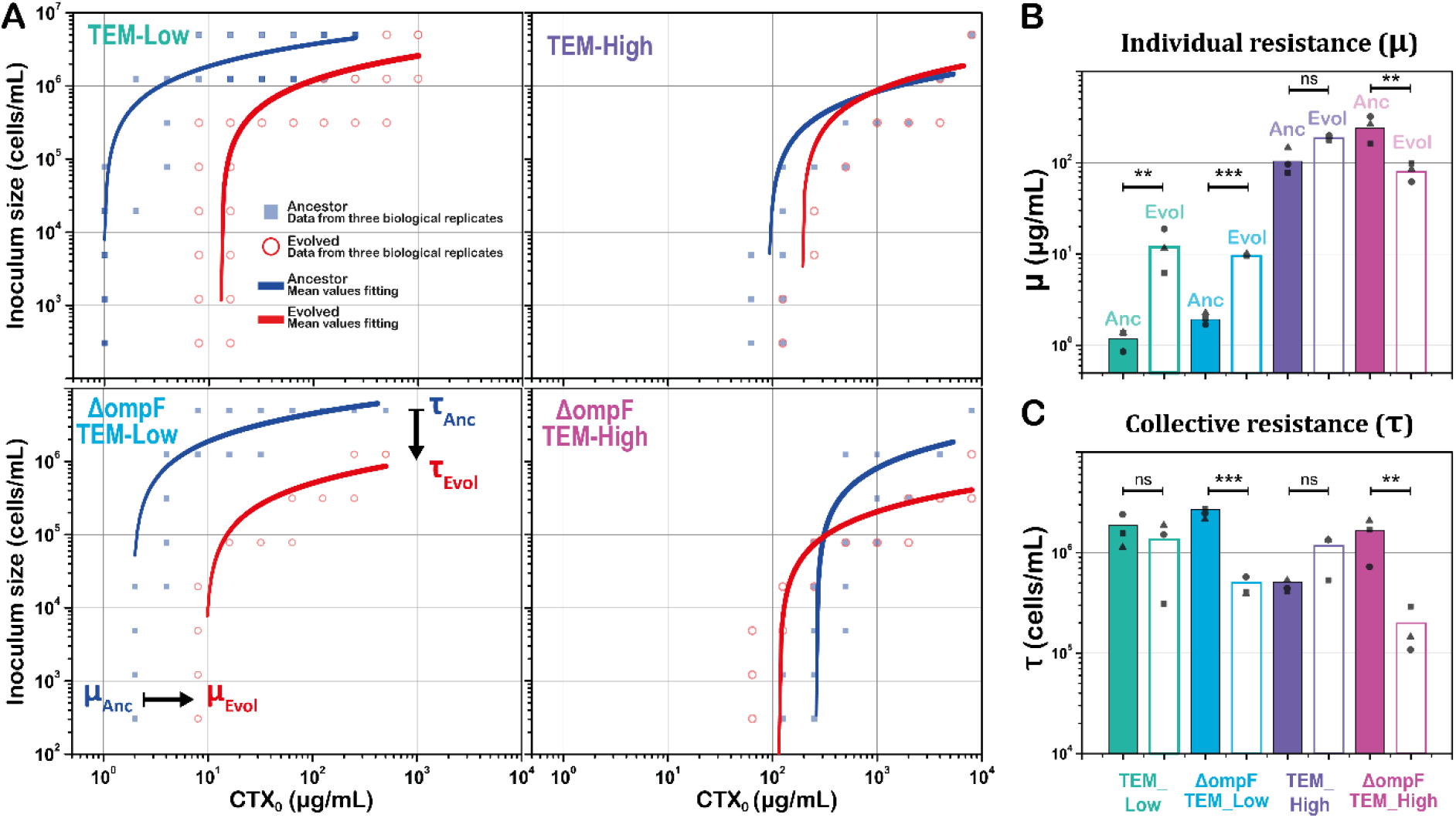
(**A)** Minimum surviving bacterial inoculum size as a function of cefotaxime concentration (CTX_0_) for ancestral strains (blue squares) and evolved lineages (red circles) derived from each of the four ancestral strains. In all cases, data from three biological replicates are shown. The mean values of experimental data were fitted (blue and red lines) to the model shown in Equation 1. The fitted parameters associated with individual resistance (µ) and collective resistance (τ) are presented in panels **(B)** and **(C)**, respectively. Statistical comparisons of means values were performed using a two-sample t-test assuming equal variances (Table S3 and S4). Significance levels are indicated as follows: ns, not significant; *(p < 0.05); **(p < 0.01); ***(p < 0.001).

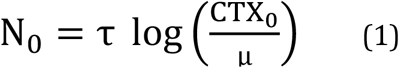

In this model, *N*_*0*_ represents the inoculum size, *CTX*_*0*_ the highest initial CTX concentration at which a given inoculum of bacteria can proliferate, *μ* approximates the single-cell MIC (scMIC) and *τ* is a parameter for the level of collectivity in CTX resistance, with smaller values of *τ* representing larger MIC increases of larger inoculum, hence greater collective CTX-resistance.

For each of the four ancestral strains (blue lines, Figure 2A) and for a random clone isolated from one of the six evolved lineages per strain (red lines, Figure 2A), we evaluated changes in μ and τ by fitting experimental measurements of the minimum surviving inoculum (MSI) obtained at different inoculum sizes as a function of the initial CTX concentration. As a general trend, strains carrying low-activity β- lactamase exhibited larger increases in individual-level resistance, showing greater changes in μ than those observed for high-activity β-lactamase producers (Figure 2B). In addition, strains with more private β-lactamase activity (ΔompF variants) displayed greater increases in collective resistance than the corresponding OmpF-producing strains (Figure 2C).

The two evolved clones carrying low-activity β-lactamase exhibited substantial increases in individual-level resistance compared with their respective ancestral strains, with μ values increasing 9-fold for TEM_Low (p = 0.003) and 5-fold for ΔompF_TEM_Low (p = 5 × 10^−5^). Differences in collective resistance were not significant for TEM_Low (p = 0.44), whereas a strong decrease in τ was observed for ΔompF_TEM_Low (p = 0.0003). Changes in individual resistance among the high-activity β- lactamase producers followed different trend. For TEM_High, a small 2-fold increase was observed, although this change was not statistically significant (p = 0.075). Surprisingly, the evolved clone of ΔompF_TEM_High showed a 3-fold decrease in μ (p = 0.01) (Figure 2B). Along with this greater CTX susceptibility at individual level, ΔompF_TEM_High showed the greatest increase of resistance at population level with 8-fold decreased in τ (p = 0.008) (Figure 2C). This observed trade-off of increased collective resistance against decreased individual-level resistance relative to the ancestral strain, suggests it evolved an altruistic form of CTX resistance.

Based on these observations, for each of the four strains, a comparison was made for all six evolved final lineages and compared with their respective ancestor using only two initial inoculum sizes: a small inoculum of 500 cells/mL and a large inoculum of 5.0×10^6^ cells/mL. Resistance level for all final evolved lineages was consistent within them and with the observations described above (Figure S5 and S6, SI)

### Whole genome sequencing analysis

To identify the molecular mechanisms underlying the observed changes in individual and collective CTX resistance, we performed whole-genome sequencing on a random clone from each of the six final evolved lineages and the corresponding ancestral strains. This revealed unique and shared mutations among the six evolved final lineages per strain, which allowed to roughly pinpoint the time mutations were fixed in each lineage. Given that the six final clones of each strain share a common history during at least the initial 17 days of the evolution experiment when the increase in CTX tolerance was greatest, we were particularly interested in the shared mutations (Figure S2 and S3, SI).

For TEM_Low lineages, two mutations were shared by all six finals, with no additional unique mutations identified (Figure S2, SI). One of those was a single nucleotide polymorphism (SNP) in the *cpxA* gene, resulting in an amino acid substitution at position 56, from glutamic acid to lysine (E56K). This gene encodes a kinase embedded in the inner membrane, and the activation of this gene has been associated with membrane stress regulation in *E. coli*, promoting increased antibiotic resistance.^44,45^ The other shared mutation is a large deletion of approximately 5 kbp mediated by an IS1-like transposable element (Table S2, SI). As annotated for REL606,^46^ the genes affected by this deletion (Table S2) encode a protein complex embedded in the inner membrane of gram-negative bacteria that is involved in the biogenesis of capsular polysaccharides associated with pathogen virulence and bacterial envelope stability.^47^

The evolved Δ*ompF*_TEM_Low clones shared a single mutation, i.e. a frameshift mutation (Δ1 bp) in the gene *rne*, encoding the enzyme RNAse E involved in mRNA maturation. In addition, two clones shared a nonsense mutation introducing a stop codon at position 248 in the gene encoding catalase KatE, involved in the regulation of oxidative stress and previously associated with resistance to cephalosporins,^48^ together with a large deletion of ∼ 11 kbp involving genes related with oxalate degradation, like *oxc* and *frc* (Table S2),^49^ which seems to be neutral in our experimental background. Another clone of Δ*ompF*_TEM_Low had a unique missense mutation (G38S) in the gene encoding BaeS, a protein associated with the reduction of oxidative stress.^50^ The six evolved TEM_High clones share a mutation (T436P) in RNAse E gene (*rne)*, together with a Δ12-bp deletion in gene *lolC*, which encodes a lipoprotein involved in the integrity of the outermembrane in *E. coli*.^45^ Additionally, two clones share missense mutation G272V in *trxB*, a gene associated with reduction of oxidative stress.^51^ Two of the final evolved lineages presented unique deletions of 0.5 and 9.5 kbp mediated by IS150 and IS1A transposable elements, respectively (Table S2). The first one is affecting the gene *rbsD*, related with uptake and metabolism of ribose as a source of carbon,^52^ which might be neutral in our experimental background. The deletion of 9.5 kbp has been observed before in similar evolution experiments and has been reported as a neutral mutation as well.^12^

However, we were most interested in understanding the observed combination of increased collective and decreased individual resistance in the evolved lineages of Δ*ompF*_TEM_High used for MSI curves presented in Figure 2A, which is highlighted in red as P4 in Figure 3. We identified two mutations in P4 that were shared among the all six evolved linages of Δ*ompF*_TEM_High (Figure 3A), both involving activity of IS1A elements. One is a 777-bp deletion caused by the excision of an IS1A element from a gene encoding outermembrane porin NmpC (Figure S7), the other a large IS1-mediated deletion of ∼ 43 kbp, corresponding to deletions observed in a similar study of experimental evolution of *E. coli* under CTX pressure developed by Schenk et al. (2022),^12^ genes involving in this big deletion are show in Table S2. In addition, P4 lineage harbored a missense mutation (G1068A) in *rpoB*, encoding the σ factor of the RNA polymerase that can regulate the transcription of genes involved in bacterial envelope biogenesis.^53^ Mutations in *rpoB* were also found previously by Schenk et al. (2022).^12^ Unique mutations in the other clones were found including another missense mutation (E13A) in *rne*, as well as missense mutations in the genes *ackA* and *lrhA*. The *ackA* encodes and acetate kinase involved in the pathway for antibiotic stress regulation, including β-lactams, mediated by the *cpxA* pathway.^44^ And *lrhA* has been pointed to establish an epistatic interaction with *rpoS* as a transcriptional regulator that can increase the expression of porins like *ompF* in *E. coli*.^54^

**Figure 3.**
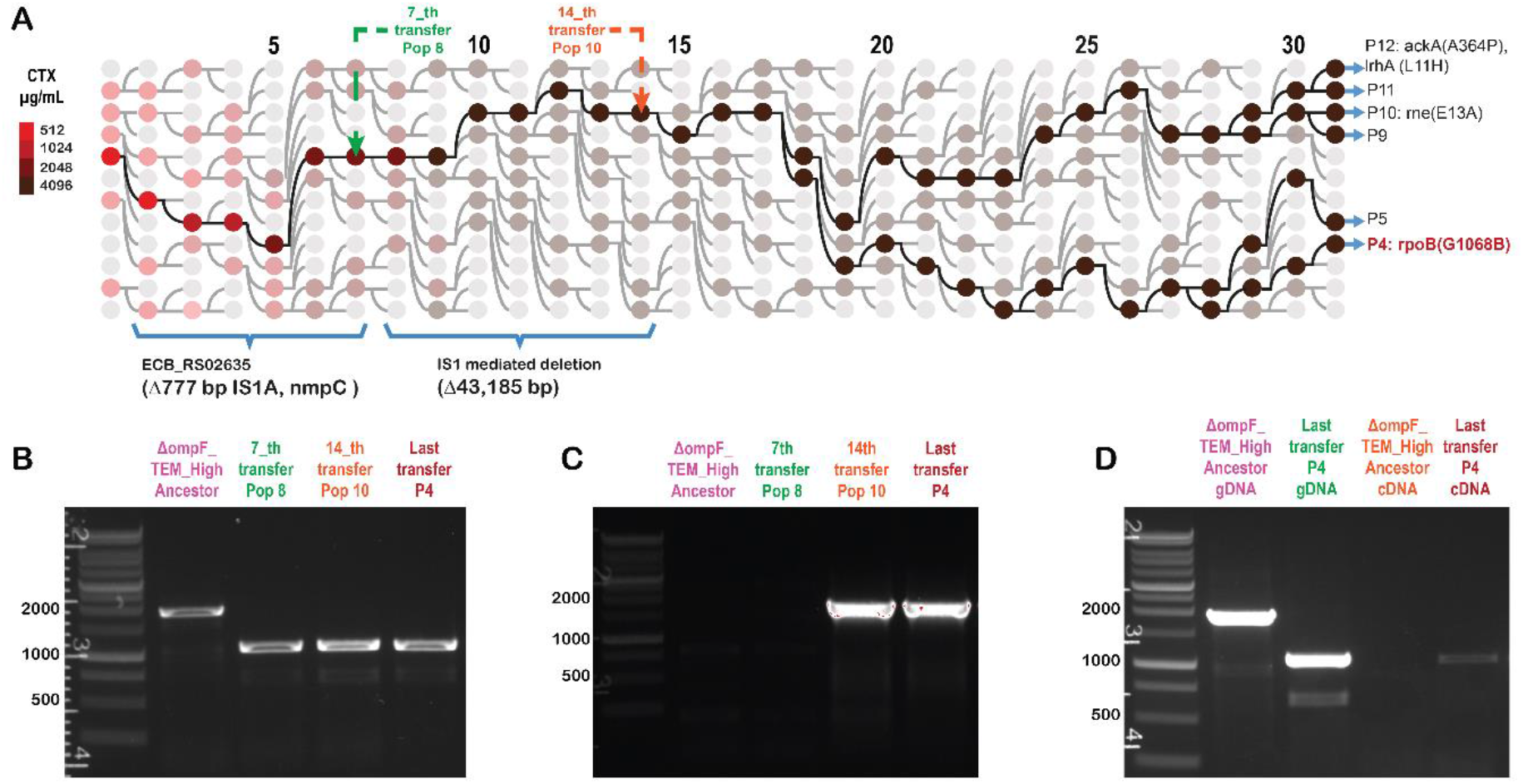
Activation of dormant porin NmpC is the first mutation in the evolved Δ*ompF*_TEM_High strains. (A) Mutations identified in the six evolved clones of strain Δ*ompF*_TEM_High. The six evolved clones shared two mutations, excision of a 777-bp IS1A from *nmpC* and a IS1-mediated 43-kbp deletion, in addition to mutations unique to some clones (shown on the right). (B) PCR amplification of the IS1A excision from *nmpC* in weekly frozen samples after 7 and 14 transfers from populations of the ancestral strain of Δ*ompF*_TEM_High, the 777-bp excision show its fixation by the 7th transfer. (C) The 44,185 bp deletion was fixed somewhere between the 7th and 14th transfers. (D) RT-PCR showing the expression of *nmpC* for the final selected lineage, marked as P4 in (A).

To better understand the role of the shared 777-bp and 43-kbp deletions, we first determined the order in which they were selected using PCR with mutation-specific primers on population samples that had been frozen each week. This revealed that the 777-bp deletion was fixed within the first seven days, while the 43-kbp deletion appeared a week later (Figure 3B and C). Therefore, our primary focus was to understand the role of the 777-bp deletion from *nmpC*. Because the 777-bp deletion generated the excision of IS1A, this mutation could have activated the expression of porin NmpC, we tested its expression by performed RT-PCR in the ancestor and a clone from P4 in Figure 3A. The primers showed the predicted bands using genomic DNA, while they showed only a product for the cDNA obtained from mRNA for the evolved clone and not from the Δ*ompF*_TEM_High ancestor (Figure 3D), suggesting the IS1A excision activated dormant outermembrane porin NmpC.

### Role of NmpC activation in individual and collective CTX resistance

To better understand how fixed mutations modify the growth dynamics of ΔompF_TEM_High, we conducted further phenotypic analyses across the lineage landscape highlighted in Figure 3A. We include the ΔompF_TEM_High ancestral line, from now on called *Ancestor*; the derived lineage after 7^th^ transfer in which only *nmpC* was activated after seven transfers; and the final evolved lineage, the triple mutant marked as P4 in Figure 3A, which in addition to *nmpC* activation, harbors the big deletion of 43,184 bp and an SNP in *rpoB*.

First, we monitored the optical density at 600 nm (OD600) over time for the three members of this landscape. Growth kinetics were assessed at both high (HI = 5 × 10^6^ cells/mL) and low (LI = 5 × 10^2^ cells/mL) inoculum sizes, using increasing CTX concentrations in each case, as shown in Figure 4, S8 and S9. We did not observe significant differences of bacterial growth in the absence of the antibiotic. And, in agreement with the observation shown in Figure 2, the resistance level of evolved lineages is higher at HI and lower at LI when compare with the Ancestor (Figure 4A, S8 and S9).

**Figure 4.**
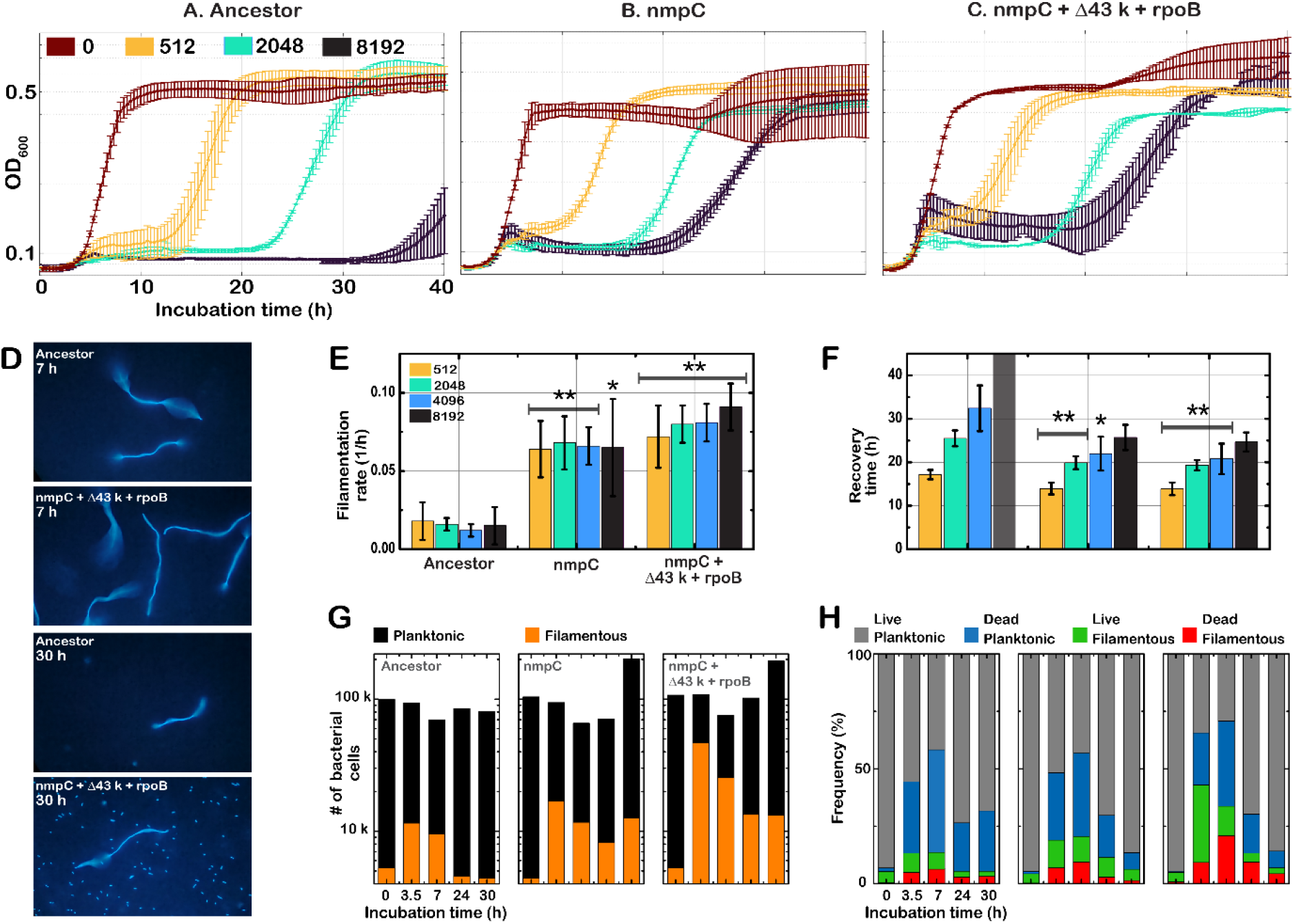
Growth curves for the three selected members the evolutionary landscape of ΔompF_TEM_High. (A to C) Optical density for each variant was monitored over 40 h, with incubation at 37 °C. Different initial CTX concentrations were used, as indicated in the labels. Curves correspond to a representative biological replicate and error bars are the standard deviation of two technical replicates, different biological replicates are shown in Figure S8 and 9. (D) Representative microscopy images showing the growth of the Ancestor strain and the triple mutant evolved lineage at different time points over the incubation period, see labels. Kinetic parameters extracted from the growth curves: (E) filamentation rate at HI (5×10^6^ cells/mL) and (F) recovery time at HI, recovery time of the Ancestor was not observed before 40 h of incubation when CTX_0_ was 8192 µg/mL. Data correspond to the mean value of three biological replicates each one composed by two technical replicates giving a total n=6. Error bars correspond to the standard deviation of the six total replicates. Significance is evaluated when comparing the Ancestor against the evolved lineage at a given CTX concentration: 0.05>p≥0.01 is * and 0.01>p≥0.001 is **, see tables S5 an S6 of supplementary file for *p*-values. (G) Total number of bacterial cells over time detected by flow cytometry in a 80 µL sample (see methods for details) indicating the fractions of planktonic and filamentous bacteria in each time point. (H) Distribution planktonic, filamentous, live and dead bacterial cells from the same samples showed in (G). Data correspond to the mean value of three technical replicates.

When bacteria were culture challenged with CTX, for all the strains at HI conditions we observed non-monotonic growth dependent on the initial concentration of CTX, as previously described,^39^ with some different trends among the strains. All the members of this landscape exhibited an initial growth phase between 0 and 7 h of incubation. However, the maximal OD reached during this first growth phase was higher for all evolved lineages than the one observed for the ancestor (Figure 4 A to C). Microscopy and flow cytometry analyses at different time points over the incubation time confirmed that this early increase in OD is associated with the production of more numerous and larger filaments of the evolved lineages than the observe for the Ancestor (Figure 4 D and G).

Further analysis of OD signal shown significant increase in the rate at which filaments grow (filamentation rate) once the *nmpC* mutation is fixed (mean values from three biological replicates, Table S5 for *p*-values). However, the filamentation rate remains constant across the different CTX concentrations evaluated and additional mutations do not produce any significant change in the filamentation rate (Figure 4E and Table S5).

At the highest CTX concentration we tested (8192 µg/mL), once the filamentation phase reaches its maximal point, the bacterial cultures undergo a decay phase in which the OD signal decreases and remains relatively low for several hours until the population begins to recover, leading to an increase in OD. As observed in Figure 4F, all mutants take significantly less time to recover from antibiotic treatment than the Ancestor (Table S5 for *p*-values). Additionally, higher initial CTX concentrations are associated with longer times for bacterial populations to reach the recovery phase. Because the lower resistance of the Ancestor strain, at the highest CTX concentration evaluated, we were unable to determine the exact recovery time during the time of the experiment. If wild-type populations are able to recover from this high CTX concentration, the process likely takes more than 40 hours. However, as shown in Figure S8, survival of Ancestor bacterial populations at 8192 µg/mL of CTX appears to be stochastic, with low repeatability among biological replicates. Finally, Additional mutation to *nmpC* do not produce significant difference in recovery time (Figure 4F and Table S5 for *p*-values).

To better understand the significant changes in bacterial growth dynamics observed at 8192 µg/mL, we carried out microscopy and flow cytometry analysis of bacteria growth over the incubation time. Microscopy observations, after 7 h of incubation clearly show a lower density of filamentous bacteria then the observed for the triple mutant. In agreement with OD measurements, after 30 hours, we did not observe any significant change in biomass by microscopy compared to that at 7 hours in the Ancestor cultures. In contrast, for the triple mutant, the density of planktonic cells increased significantly at this time point, indicating proliferation of the triple mutant population (Figure 4D).

Flow cytometry analyses were also consistent with the OD measurements and microscopy observations, while providing additional insights. First, we correlated the SSC signal with blue fluorescence intensity to distinguish planktonic from filamentous bacteria. Furthermore, dead bacteria were stained with Propidium Iodide and identified by red fluorescence, enabling their differentiation from live bacteria (Figure 4G and H).

This analysis demonstrated that all samples started with similar bacterial densities and comparable distributions of planktonic, filamentous, live, and dead cells. After 3.5 h of incubation, the fraction of filamentous bacteria increased in all strains, remaining lowest in the Ancestor strain and increasing in both the *nmpC* mutant and the triple mutant, with the latter exhibiting the highest fraction of filamentous cells. Importantly, the total number of bacteria did not change significantly after 3.5 h of incubation, indicating that the increase in OD signal was primarily due to biomass accumulation through filamentous growth rather than bacterial cell division and proliferation. Furthermore, the decrease in OD signal observed after 7 h of incubation correlated with a reduction in the total number of bacteria (Figure 4G). However, the triple mutant exhibited a significantly higher fraction of dead filamentous cells than either the Ancestor or the *nmpC* mutant (Figure 4H). This observation suggests that the two additional mutations in the triple mutant contribute to stabilizing the altruistic resistance mechanism that evolved in this landscape. After 24 h of incubation, the fraction of filamentous bacteria decreased in the Ancestor strain, without significant changes in the total number of bacteria, even after 30 h of incubation. In contrast, after 24 h the mutant lineages entered a recovery phase characterized by an increase in the total bacterial population, an increase in the fraction of live planktonic bacteria, and a reduction in the fractions of filamentous and dead cells (Figure 4H).

When cultures were initiated at a low inoculum size in the presence of CTX, we did not observe non-monotonic behavior. We hypothesize that the filamentation phase still occurs but is not detectable because it remains below the detection limit. Consistent with the observations in Figure 2, populations from the Ancestor line were able to grow at higher concentrations than populations from the evolved lines when cultures were started at 5 × 10^2^ cells/mL (Figure S9). However, under these conditions, stochasticity in bacterial growth did not yield any clear trend or significant differences (Figure S10 and Table S6). Furthermore, when we assessed growth rates in the absence of CTX, we observed no differences associated with the accumulation of mutations or the inoculum size (Supplementary Figure S10).

### CTX degradation during incubation

We used high-performance liquid chromatography (HPLC) to monitor CTX concentrations in the supernatant over time in cultures of both Ancestor and triple-mutant bacteria. Cultures at both high and low inoculum sizes were monitored using initial CTX concentrations of 4096 µg/mL and 64 µg/mL, respectively (Figure 5 and S11 for extended data).

**Figure 5.**
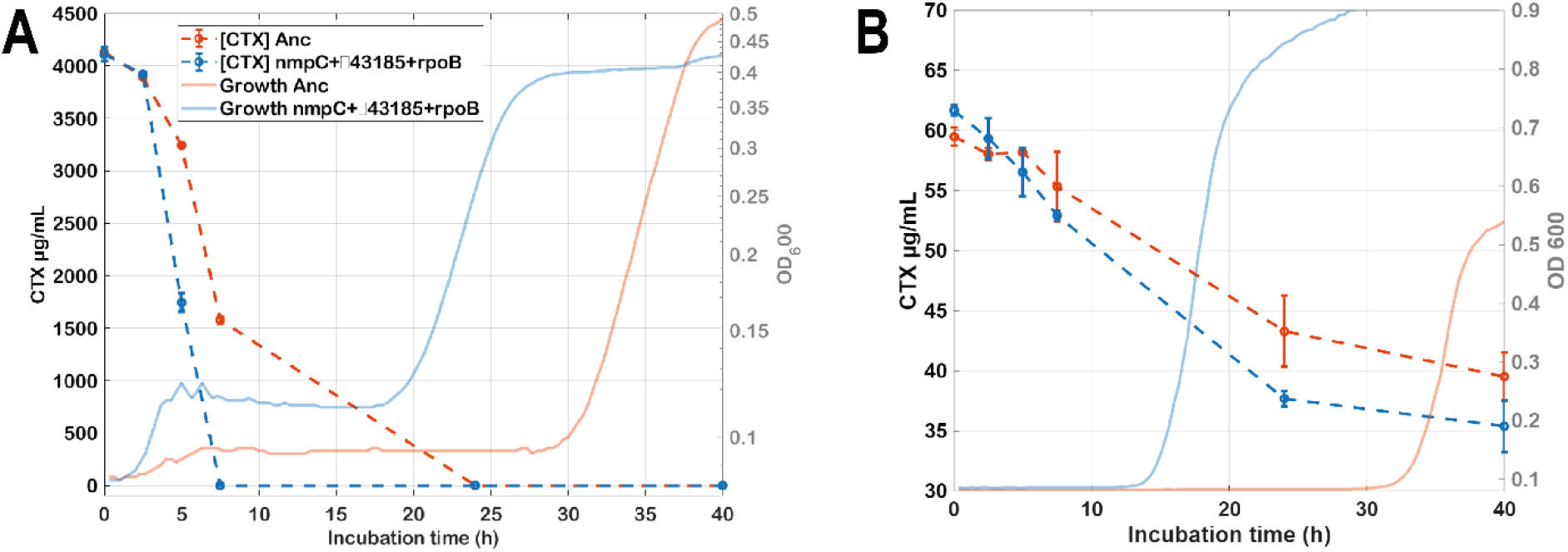
(A) Degradation rate of CTX over time in bacterial cultures of the ΔompF_TEM_High and its evolved lineage, the triple mutant (nmpC + Δ43,184 bp + rpoB). Cultures were initiated at 5×10^6^ cells/mL and 4096 µg/mL of CTX. (B) Cultures were started at 5×10^2^ cells/mL and 64 µg/mL of CTX. In both cases the left axis shows CTX concentration over incubation time. Each data point represents the mean of two technical replicates, and error bars indicate the standard deviation. The right axis shows the growth curves recorded for the bacterial cultures from which samples were taken for HPLC analysis.

For the high-inoculum cultures, CTX concentrations in the supernatant remained nearly identical in Ancestor and triple-mutant cultures during the first 2.5 h of incubation. However, after this time point, CTX levels decreased more rapidly. In the triple-mutant culture, CTX dropped below the detection limit shortly after 7 h of incubation, approximately coinciding with the time at which the initial filamentation phase reached its maximum signal. In contrast, ancestor cultures maintained detectable CTX levels for more than 7 h, demonstrating a faster degradation rate in the triple-mutant compared to the ancestor bacterial cultures (Figure 5A). In the low-inoculum cultures, CTX degradation took longer than 7 h. In these cultures, CTX remained detectable in both the Ancestor and the evolved lineage even at the final time point measured (40 h of incubation). However, the evolved lineage again degraded CTX faster than the Ancestor strain (Figure 5B).

## Discussion

Among the four evolved ancestral strains, those with lower initial permeability exhibited a greater collective potential for resistance, as reflected by lower τ values (Figure 3). As expected, relatively low permeability provides the potential to evolve collective resistance mechanisms. However, an unexpected and interesting pattern emerged when low initial permeability was combined with high catalytic efficiency. This combination produced a trade-off in the antibiotic resistance of evolved lineages derived from the ΔompF_TEM_High ancestral strain. These evolved lineages exhibited higher MIC values than their ancestor at inoculum densities above 1 × 10^3^ cells/mL, whereas the opposite trend was observed at inoculum densities below this threshold (Figure 2). Together with this phenotypic observation, and given that all six final mutants (Figures S5 and S6) as well as the single mutant harboring *nmpC* showed the same trade-off (Figures S8 and S9), we propose that *nmpC* activation is the mutation driving this altruistic behavior. And, the additional fixed mutations (including the 43,184-bp deletion and the unique mutations present in the final lineages) likely contribute to stabilizing the observed altruistic resistance mechanism.

*NmpC* activation became fixed early during the evolution experiment, occurring no later than after seven transfers (Figure 3). Its activation led to a significant increase in the initial filamentation rate and a reduction in the recovery time of bacterial cultures challenged with CTX. Microscopy and flow cytometry analyses revealed that evolved strains formed longer filaments that produced more biomass during the first 5 hours of incubation than the ΔompF_TEM_High ancestral strain. In addition, during the first 7 hours of incubation, the proportion of dead filaments increased in the evolved lineages, reaching its highest level in the triple mutant (Figure 4H). However, despite the greater increase in dead filaments observed in the triple mutant, no major differences in recovery time were detected compared with the single mutant (*nmpC*). Furthermore, the growth rates for the ancestor and evolved lineages in in the absence of CTX were not significantly different (Figure S10), suggesting that *nmpC* activation provides a benefit primarily in environments containing CTX. Finally, HPLC measurements showed that CTX concentrations decreased rapidly during the first 10 h of incubation, with faster degradation observed in populations of the triple mutant evolved lineage.

Besed on result summarized in the previous paragraph, N*mpC* activation may increase the permeability to CTX molecules, thereby increasing the effective periplasmic concentration in mutants with activated *nmpC*. As a result, because the periplasmic concentration and efficiency of β-lactamase are not sufficient to immediately degrade all incoming CTX molecules, some CTX molecules bind to PBP3, its primary target. The increased inhibition of PBP3 leads to longer filaments in *nmpC* mutants than in the Ancestor. Hence, while bacteria grow as filaments, intracellular periplasmic degradation may be more efficient than the one happening in planktonic bacteria. Thus, in mutants with higher permeability, more antibiotic molecules are in the periplasm space increasing the likely of interaction with β- lactamases and the rate at which CTX is degraded indirectly reducing extracellular environmental CTX concentration as a weak altruistic collective resistance.

Because of their lower permeability, wild-type bacteria are slower to reduce antibiotic concentrations. Consequently, when the initial CTX concentration exceeds 4096 µg/mL, the last concentration at which populations of these bacteria can survive, so the ancestor population goes extinct before the medium is fully detoxified. In contrast, faster degradation in *nmpC* mutants enables more rapid detoxification of the medium, allowing these populations to survive at concentrations higher than 4096 µg/mL, as was shown in Figure 4.

At low inoculum, the faster intracellular degradation associated with increased permeability does not lead to the same population-level response. Higher CTX permeability increased the likelihood of cell death, as is show in Figure 4H. Therefore, although *nmpC* mutants degrade CTX faster, they also die faster than the ancestor when CTX is present. As a result, if the initial population is not large enough, the *nmpC* mutant population goes extinct before the populations of the ancestor if they start at the same low bacterial density,^19^ explaining the population-dependent trade-off observed in Figure 2.

Our observations strongly support that the increased resistance in evolved lineages derived from ΔompF_TEM-High is driven by enhanced intracellular degradation. However, extracellular degradation resulting from cell lysis may also contribute to the overall resistance level. At present, there is insufficient evidence to fully support this hypothesis. Demonstrating a higher death rate in the presence of the antibiotic, together with increased extracellular enzymatic degradation, would be necessary to establish that both intracellular degradation within filaments and enzymatic extracellular degradation act synergistically in CTX resistance mechanisms.

### Materials and Methods. Bacteria strains and media

Four different ancestral strains were used in this study (Table S1, Supplementary Information), all derived from *E. coli* B REL606 and modified to express Blue Fluorescent Protein (BFP) through the insertion of cat-J23101-mTagBFP2 into the galK gene. To obtain the TEM-Low and TEM-High variants, electrocompetent cells derived from a single colony were transformed with the pACTEM plasmid, introducing a single mutation into the original TEM-1 sequence, as show in Table S1 of the Supplementary Information. Finally, the ompF gene was deleted using the Red/ET recombination protocol from Gene Bridges (Heidelberg, Germany), generating the ΔompF-TEM_Low and ΔompF-TEM_High variants.^12,55^

All experiments were conducted using M9 medium supplemented with IPTG, glucose and casamino acids (6.8 g/L Na_2_HPO_4_·2H_2_O, 3 g/L KH_2_PO_4_, 0.5 g/L NaCl, 1 g/L NH_4_Cl, 1 mM MgSO_4_·7H2O, 0.1 mM CaCl_2_, 0.2% casamino acids, and 0.4% glucose, 50µM IPTG). When required, 1.5% (w/v) agar was added to obtain solid medium.

### EVOLUTION experiment

Each ancestral line was independently cultured on agar plates for 24 h at 37 °C using half of the corresponding MIC of cefotaxime (CTX; Duchefa). A different colony was then used to initiate twelve independent subpopulations from each ancestral line in M9 liquid medium. Each population was exposed to four consecutive initial CTX concentrations: 0.25, 0.5, 1, and 2 times the MIC of the corresponding ancestral line. After 24 h of incubation at 37 °C, the lineages showing the highest bacterial growth, as determined by optical density measurements (BioTek Synergy H1, Agilent), at the highest CTX concentration that still permitted growth were selected.

One percent of the culture volume (2 µL) from each selected population was transferred to two columns of a new 96-well plate containing fresh medium and four new CTX concentrations. The minimum CTX concentration in the new plate corresponded to the highest concentration at which most of the selected population showed growth in the previous round, while the other three concentrations were obtained by progressively doubling this minimum concentration. This procedure was repeated for 30 rounds, allowing the strains to evolve over approximately 200 generations (Supplementary Figure S1, SI).

### Minimal survival inoculum determination

Based on the protocol described by Geyrhofer et al.,^19^ MSI curves were determined for each ancestral line and one evolved lineage from each ancestral line. Briefly, the population size was varied across the eight rows of a 96-well plate, starting with row A at the highest inoculum (5×10^6^ cells/mL) and proceeding with 4-fold serial dilutions down to row H, which contained the lowest inoculum used (3.05×10^2^ cells/mL). CTX concentrations varied across the 12 columns of the plate, following 2-fold serial dilution steps. The CTX concentration range used for the TEM_Low ancestral line was 0.512 to 2.5×10^-4^ mg/mL; for the TEM_Low evolved lineage and the ΔompF_TEM_Low ancestral line, it was 1.0 to 5×10^-4^ mg/mL; and for the ΔompF_TEM_Low evolved lineage, it was 4.096 to 2×10^-3^ mg/mL. Finally, for all TEM_High variants, with and without ompF, the CTX concentration gradient ranged from 16 to 8×10^-3^ mg/mL. MSI Curves were evaluated after 40 h of incubation at 37 °C. The initial antibiotic concentration in the last well showing detectable growth was considered the MSI value for that CTX concentration and one point in the whole MSI curve (Figure S4, Supplementary Information). Experimental data was plotted and fitted to the model shown in **eq 1** using the *cftool* in MATLAB (R2025b).

### Whole genome characterization

After 30 serial transfers during the evolution experiment, each of the six final evolved populations generated from each ancestor, as well as its respective ancestral population (Supplementary Figures S2 and S3, SI), was independently plated on M9 agar. For the ancestral lines, M9 agar plates were supplemented with CTX at concentrations corresponding to 0.5× the previously determined MIC, and for evolved lineages 1× the MIC of its corresponding ancestral line (Supplementary Table S1, SI). After 40 h of incubation at 37 °C, one colony was picked from each culture plate and their total genomic DNA was isolated using the DNeasy Blood and Tissue Kits for DNA Isolation (QIAGEN), manufacturer’s instructions were followed.

The Nextera XT (Illumina) kit was used for library preparation, following the in-house protocol known as *hackFlex*. Libraries were sequenced on a HiSeq 2500 using paired-end 150-bp (PE150) sequencing through the sequencing service provided by Novogene.

Ancestral and evolved whole genomes were analyzed using the *Breseq* pipeline,^56^ with REL606 (taxonomy ID: 413997, NC_012967.1, NCBI) and pacTEM1 (taxonomy ID:2724077, MN386081.1, GenBank) as the reference for chromosomal genome and plasmid, respectively. Mutations shared between the ancestral lines and their corresponding evolved lineages were filtered out, retaining only mutations that were present at the evolved lineages after 30 serial transfers.

For the PCR reactions, DNA templates were prepared from a single colony for each sample by resuspending the colony in 50 µL of DNase-free water and heating it for 10 min at 90 °C. PCR reactions were performed following a standard protocol using GoTaq polymerase. The primers used to confirm the 777-bp deletion were: forward, 3′-TTAGAACTGATACACCAGACCT-5′, and reverse, 3′- ATGAAAAAATTAACAGTGGCA-5′. The primers used to confirm the IS1-mediated 43-kbp deletion were: forward, 3′-TTCCGAGAATGGACACCAGC-5′, and reverse, 3′- GCCACAAGCAATACCGACAG-5′.

NmpC expression was assessed using RT-PCR. RNA was isolated using the RNeasy Mini Kit (QIAGEN), and cDNA was synthesized using the SensiFAST™ cDNA Synthesis Kit (Bioline), according to the manufacturers’ instructions. The same primers used to confirm the 777-bp deletion were used to assess NmpC expression. PCR products were separated by electrophoresis on 1.0% agarose gels, stained with ethidium bromide, and visualized using a gel documentation system (Gel Doc XR+, Lab^TM^ Software, Bio-Rad).

### Growth kinetics

Growth kinetics of selected lineages were monitored for 40 h in a 96-well plate format using a plate reader (Synergy H1M2, Agilent BioTek). Optical density at 600 nm (OD_600_) was measured every 15 min. The temperature was maintained at 37 °C, and the plate was shaken for 2 min before each OD_600_ measurement. Filamentation growth rate, dead rate, recovery time and recovery rate parameters were extracted from growth curves using a home-made code in MATLAB (R2025b). When required, a replicate plate was incubated in an oven, and samples were collected from it every 2.5 h for microscopy, HPLC, or flow cytometry analyses.

Fluorescence microscopy images were acquired using an Axiophot microscope (ZEISS) equipped with a light source (X-Cite Series 120, EXFO). Five-microliter samples were placed between a glass slide and a coverslip and observed under oil immersion using a 63× objective (63×/1.4, ZEISS). Images were captured using a smartphone camera attached to the microscope binoculars.

Flow cytometry was carried out using a MACSQuant system. Ten-microliter samples were taken from bacteria growing in wells and diluted 10 times using M9 liquid media and then analyzed by flow cytometry. SSC threshold was fixed a 1.4 AU and samples were always running at hits rate around 1000 events/s. Based on the signal obtained for M9-only and bacterial suspension at different time point we set the gating strategy illustrated in the Supplementary Figure S11 (SI).

### HPLC analysis

At each time point, the entire volume from one technical replicate was collected from the corresponding well and immediately centrifuged for 3 min at 800 rpm. The pellet was discarded, and the supernatant was filtered through a 0.2 µm pore-size filter. Two microliters of 10 mM clavulanic acid solution was then added to the supernatant to inhibit β-lactamase activity. The samples were subsequently frozen at −20 °C and stored until all time points had been collected. Once all samples had been collected, they were slowly thawed on ice and analyzed in a single batch using an Ultra-High-Performance Liquid Chromatography (UHPLC) system (UltiMate™ 3000, Thermo Scientific), equipped with a LiChrospher® 100 RP-18 C18 reversed-phase column (125 mm × 4 mm, 5 µm; Merck). Two technical replicates were analyzed for each bacterial strain.

The mobile phase consisted of ultrapure water containing 0.2 mM H_3_PO_4_ (A) and methanol (B). Samples were analyzed using an isocratic method with a 70:30 A/B (v/v) ratio for 20 min at a flow rate of 1.0 mL/min. Samples were injected at a volume of 10 µL, and compounds were detected by UV absorbance at 310 nm. Data were recorded and analyzed using Chromeleon™ software (v. 7.2.10, Thermo Scientific).

Cefotaxime (CTX) peaks were identified by comparing their retention time with that of a CTX standard solution. A calibration curve was generated using the same standard solution by preparing six consecutive twofold serial dilutions, starting at 4096 µg/mL. The peak area was plotted against the corresponding CTX concentration to generate a calibration curve, which was subsequently used to determine the CTX concentration in samples collected from growing bacterial cultures.

## Supporting information

SI_Ochoa_2026

## Acknowledgements

A.O. thanks the Secretariat of Education, Science, Technology and Innovation of Mexico City (SECTEI) for the fellowship granted under Agreement No. SECTEI/069/2024, and the Genetics Group at Wageningen University & Research for hosting agreement established to support this project.

## References

(1) World Health Organization. Global Antibiotic Resistance Surveillance Report 2025. World Health Organization; 2025. doi:10.2471/B09585.

(2) Murray CJL, Ikuta KS, Sharara F, et al. Global Burden of Bacterial Antimicrobial Resistance in 2019: A Systematic Analysis. The Lancet 2022, 399 (10325), 629–655. 10.1016/S0140-6736(21)02724-0.

(3) Klein, E. Y.; Impalli, I.; Poleon, S.; Denoel, P.; Cipriano, M.; Van Boeckel, T. P.; Pecetta, S.; Bloom, D. E.; Nandi, A. Global Trends in Antibiotic Consumption during 2016-2023 and Future Projections through 2030. Proc. Natl. Acad. Sci. U. S. A. 2024, 121 (49). 10.1073/pnas.2411919121.

(4) Mora-Ochomogo, M.; Lohans, C. T. β-Lactam Antibiotic Targets and Resistance Mechanisms: From Covalent Inhibitors to Substrates. RSC Medicinal Chemistry. Royal Society of Chemistry October 1, 2021, pp 1623–1639. 10.1039/d1md00200g.

(5) Montaner, M.; Lopez-Argüello, S.; Oliver, A.; Moya, B. PBP Target Profiling by β-Lactam and β-Lactamase Inhibitors in Intact Pseudomonas Aeruginosa: Effects of the Intrinsic and Acquired Resistance Determinants on the Periplasmic Drug Availability. Microbiol. Spectr. 2023, 11 (1). 10.1128/spectrum.03038-22.

(6) Bertsche, U.; Kast, T.; Wolf, B.; Fraipont, C.; Aarsman, M. E. G.; Kannenberg, K.; Von Rechenberg, M.; Nguyen-Distèche, M.; Den Blaauwen, T.; Höltje, J. V.; Vollmer, W. Interaction between Two Murein (Peptidoglycan) Synthases, PBP3 and PBP1B, in Escherichia Coli. Mol. Microbiol. 2006, 61 (3), 675–690. 10.1111/j.1365-2958.2006.05280.x.

(7) Käshammer, L.; van den Ent, F.; Jeffery, M.; Jean, N. L.; Hale, V. L.; Löwe, J. Cryo-EM Structure of the Bacterial Divisome Core Complex and Antibiotic Target FtsWIQBL. Nat. Microbiol. 2023, 8 (6), 1149–1159. 10.1038/s41564-023-01368-0.

(8) Buijs, J.; Dofferhoff, A. S. M.; Mouton, J. W.; Wagenvoort, J. H. T.; Van Der Meer, J. W. M. Concentration-Dependency of β-Lactam-Induced Filament Formation in Gram-Negative Bacteria. Clinical Microbiology and Infection 2008, 14 (4), 344–349. 10.1111/j.1469-0691.2007.01940.x.

(9) Aguilar-Luviano, O. B.; Santos-Escobar, F.; Orozco-Barrera, S.; Peña-Miller, R. Conditional Filamentation Enhances Bacterial Survival in Toxic Environments. May 13, 2025. 10.1101/2025.05.13.653778.

(10) Kjeldsen, T. S. B.; Sommer, M. O. A.; Olsen, J. E. Extended Spectrum β-Lactamase-Producing Escherichia Coli Forms Filaments as an Initial Response to Cefotaxime Treatment. BMC Microbiol. 2015, 15 (1). 10.1186/s12866-015-0399-3.

(11) Hussain, H. I.; Aqib, A. I.; Seleem, M. N.; Shabbir, M. A.; Hao, H.; Iqbal, Z.; Kulyar, M. F. e. A.; Zaheer, T.; Li, K. Genetic Basis of Molecular Mechanisms in β-Lactam Resistant Gram-Negative Bacteria. Microb. Pathog. 2021, 158. 10.1016/j.micpath.2021.105040.

(12) Schenk, M. F.; Zwart, M. P.; Hwang, S.; Ruelens, P.; Severing, E.; Krug, J.; de Visser, J. A. G. M. Population Size Mediates the Contribution of High-Rate and Large-Benefit Mutations to Parallel Evolution. Nat. Ecol. Evol. 2022, 6 (4), 439–447. 10.1038/s41559-022-01669-3.

(13) Bush, K.; Bradford, P. A. β-Lactams and β-Lactamase Inhibitors: An Overview. Cold Spring Harb. Perspect. Med. 2016, 6 (8). 10.1101/cshperspect.a025247.

(14) Bush, K. Bench-to-Bedside Review: The Role of β-Lactamases in Antibiotic-Resistant Gram-Negative Infections. Crit. Care 2010, 14 (224), 1–8.

(15) Du, D.; Wang-Kan, X.; Neuberger, A.; van Veen, H. W.; Pos, K. M.; Piddock, L. J. V.; Luisi, B. F. Multidrug Efflux Pumps: Structure, Function and Regulation. Nature Reviews Microbiology. Nature Publishing Group September 1, 2018, pp 523–539. 10.1038/s41579-018-0048-6.

(16) Coldham, N. G.; Webber, M.; Woodward, M. J.; Piddock, L. J. V. A 96-Well Plate Fluorescence Assay for Assessment of Cellular Permeability and Active Efflux in Salmonella Enterica Serovar Typhimurium and Escherichia Coli. Journal of Antimicrobial Chemotherapy 2010, 65 (8), 1655–1663. 10.1093/jac/dkq169.

(17) Maher, C.; Hassan, K. A. The Gram-Negative Permeability Barrier: Tipping the Balance of the in and the Out. mBio. American Society for Microbiology December 1, 2023. 10.1128/mbio.01205-23.

(18) Sugawara, E.; Nikaido, H. OmpA Is the Principal Nonspecific Slow Porin of Acinetobacter Baumannii. J. Bacteriol. 2012, 194 (15), 4089–4096. 10.1128/JB.00435-12.

(19) Geyrhofer, L.; Ruelens, P.; Farr, A. D.; Pesce, D.; de Visser, J. A. G. M.; Brenner, N. Minimal Surviving Inoculum in Collective Antibiotic Resistance. mBio 2023, 14 (2). 10.1128/mbio.02456-22.

(20) Cartagena, A. J.; Taylor, K. L.; Lopez, L. C.; Su, J.; Smith, J. T.; Manson, A. L.; Chen, J. D.; Pierce, V. M.; Earl, A. M.; Bhattacharyya, R. P. The Carbapenem Inoculum Effect Provides Insights into the Molecular Mechanisms Underlying Carbapenem Resistance in the Enterobacterales. mBio 2025, 16 (11), 1–20. 10.1128/mbio.01540-25.

(21) Vega, N. M.; Gore, J. Collective Antibiotic Resistance: Mechanisms and Implications. Current Opinion in Microbiology. Elsevier Ltd October 1, 2014, pp 28–34. 10.1016/j.mib.2014.09.003.

(22) Saebelfeld, M.; Das, S. G.; Hagenbeek, A.; Krug, J.; De Visser, J. A. G. M. Stochastic Establishment of β-Lactam-Resistant Escherichia Coli Mutants Reveals Conditions for Collective Resistance. Proceedings of the Royal Society B: Biological Sciences 2022, 289 (1974). 10.1098/rspb.2021.2486.

(23) Artemova, T.; Gerardin, Y.; Dudley, C.; Vega, N. M.; Gore, J. Isolated Cell Behavior Drives the Evolution of Antibiotic Resistance. Mol. Syst. Biol. 2015, 11 (7). 10.15252/msb.20145888.

(24) Salas, J. R.; Jaberi-Douraki, M.; Wen, X.; Volkova, V. V. Mathematical Modeling of the “Inoculum Effect”: Six Applicable Models and the MIC Advancement Point Concept. FEMS Microbiol. Lett. 2020, 367 (5). 10.1093/femsle/fnaa012.

(25) Tan, C.; Phillip Smith, R.; Srimani, J. K.; Riccione, K. A.; Prasada, S.; Kuehn, M.; You, L. The Inoculum Effect and Band-Pass Bacterial Response to Periodic Antibiotic Treatment. Mol. Syst. Biol. 2012, 8. 10.1038/msb.2012.49.

(26) Sorg, R. A.; Lin, L.; van Doorn, G. S.; Sorg, M.; Olson, J.; Nizet, V.; Veening, J. W. Collective Resistance in Microbial Communities by Intracellular Antibiotic Deactivation. PLoS Biol. 2016, 14 (12). 10.1371/journal.pbio.2000631.

(27) Iredell, J.; Brown, J.; Tagg, K. Antibiotic Resistance in Enterobacteriaceae: Mechanisms and Clinical Implications. BMJ (Online). BMJ Publishing Group February 8, 2016. 10.1136/bmj.h6420.

(28) Brepoels, P.; De Wit, G.; Lories, B.; Belpaire, T. E. R.; Steenackers, H. P. Selective Pressures for Public Antibiotic Resistance. Critical Reviews in Microbiology. Taylor and Francis Ltd. 2024. 10.1080/1040841X.2024.2367666.

(29) Yurtsev, E. A.; Conwill, A.; Gore, J. Oscillatory Dynamics in a Bacterial Cross-Protection Mutualism. Proc. Natl. Acad. Sci. U. S. A. 2016, 113 (22), 6236–6241. 10.1073/pnas.1523317113.

(30) Zhao, X.; Ruelens, P.; Farr, A. D.; de Visser, J. A. G. M.; Baraban, L. Population Dynamics of Cross-Protection against β-Lactam Antibiotics in Droplet Microreactors. Front. Microbiol. 2023, 14. 10.3389/fmicb.2023.1294790.

(31) Wang, Q.; Wei, S.; Madsen, J. S. Cooperative Resistance Varies among β-Lactamases in E. Coli, with Some Enabling Cross-Protection and Sustained Extracellular Activity. Commun. Biol. 2025, 8 (1), 968. 10.1038/s42003-025-08392-2.

(32) Frost, I.; Smith, W. P. J.; Mitri, S.; Millan, A. S.; Davit, Y.; Osborne, J. M.; Pitt-Francis, J. M.; MacLean, R. C.; Foster, K. R. Cooperation, Competition and Antibiotic Resistance in Bacterial Colonies. ISME Journal 2018, 12 (6), 1582–1593. 10.1038/s41396-018-0090-4.

(33) Mora-Ochomogo, M.; Jeffs, M. A.; Liu, J. L.; Lohans, C. T. Contributions of β-Lactamase Substrate Specificity and Outer Membrane Permeability to the Antibiotic Sheltering of β-Lactam-Susceptible Bacteria. April 22, 2025. 10.1101/2025.04.22.649001.

(34) Pagès, J. M.; James, C. E.; Winterhalter, M. The Porin and the Permeating Antibiotic: A Selective Diffusion Barrier in Gram-Negative Bacteria. Nature Reviews Microbiology. Nature Publishing Group 2008, pp 893–903. 10.1038/nrmicro1994.

(35) Delcour, A. H. Outer Membrane Permeability and Antibiotic Resistance. Biochimica et Biophysica Acta - Proteins and Proteomics. May 2009, pp 808–816. 10.1016/j.bbapap.2008.11.005.

(36) Matsumura, N.; Minami, S.; Araki, H.; Hori, R.; Ogake, N.; Watanabe, Y.; Matsumura, N.; Minami, S.; Araki, H Hori, · R; Ogake, · N; Watanabe Y. Determination of Intracellular and Extracellular-Lactamase Activities of Pseudomonas Aeruginosa after Exposure to-Lactams in Vitro and in Vivo; 2000; Vol. 6.

(37) Kim, S. W.; Park, S. Bin; Im, S. P.; Lee, J. S.; Jung, J. W.; Gong, T. W.; Lazarte, J. M. S.; Kim, J.; Seo, J. S.; Kim, J. H.; Song, J. W.; Jung, H. S.; Kim, G. J.; Lee, Y. J.; Lim, S. K.; Jung, T. S. Outer Membrane Vesicles from β-Lactam-Resistant Escherichia Coli Enable the Survival of β-Lactam-Susceptible E. Coli in the Presence of β-Lactam Antibiotics. Sci. Rep. 2018, 8 (1). 10.1038/s41598-018-23656-0.

(38) Tanouchi, Y.; Pai, A.; Buchler, N. E.; You, L. Programming Stress-induced Altruistic Death in Engineered Bacteria. Mol. Syst. Biol. 2012, 8 (1). 10.1038/msb.2012.57.

(39) Gross, R.; Mungan, M.; Das, S. G.; Yüksel, M.; Bollenbach, T.; Krug, J.; Arjan de Visser, J. G. Collective β-Lactam Resistance in Escherichia Coli Due to β-Lactamase Release upon Cell Death; 2024.

(40) Meredith, H. R.; Andreani, V.; Ma, H. R.; Lopatkin, A. J.; Lee, A. J.; Anderson, D. J.; Batt, G.; You, L. M I C R O B I O L O G Y Applying Ecological Resistance and Resilience to Dissect Bacterial Antibiotic Responses; 2018.

(41) Wen, L.; Bai, Y.; Lan, Y.; Shen, Y.; She, X.; Dong, P.; Wang, T.; Fu, X.; Huang, S. Strong Segregation Promotes Self-Destructive Cooperation. ISME J. 2025, 19 (1). 10.1093/ismejo/wraf043.

(42) Durand, P. M.; Sym, S.; Michod, R. E. Programmed Cell Death and Complexity in Microbial Systems. Current Biology. Cell Press July 11, 2016, pp R587–R593. 10.1016/j.cub.2016.05.057.

(43) Nedelcu, A. M.; Driscoll, W. W.; Durand, P. M.; Herron, M. D.; Rashidi, A. On the Paradigm of Altruistic Suicide in the Unicellular World. Evolution (N. Y). 2011, 65 (1), 3–20. 10.1111/j.1558-5646.2010.01103.x.

(44) Jing, W.; Liu, J.; Wu, S.; Li, X.; Liu, Y. Role of CpxA Mutations in the Resistance to Aminoglycosides and β-Lactams in Salmonella Enterica Serovar Typhimurium. Front. Microbiol. 2021, 12. 10.3389/fmicb.2021.604079.

(45) Grabowicz, M.; Silhavy, T. J. Envelope Stress Responses: An Interconnected Safety Net. Trends in Biochemical Sciences. Elsevier Ltd March 1, 2017, pp 232–242. 10.1016/j.tibs.2016.10.002.

(46) Jeong, H.; Barbe, V.; Lee, C. H.; Vallenet, D.; Yu, D. S.; Choi, S. H.; Couloux, A.; Lee, S. W.; Yoon, S. H.; Cattolico, L.; Hur, C. G.; Park, H. S.; Ségurens, B.; Kim, S. C.; Oh, T. K.; Lenski, R. E.; Studier, F. W.; Daegelen, P.; Kim, J. F. Genome Sequences of Escherichia Coli B Strains REL606 and BL21(DE3). J. Mol. Biol. 2009, 394 (4), 644–652. 10.1016/j.jmb.2009.09.052.

(47) Doyle, L.; Ovchinnikova, O. G.; Myler, K.; Mallette, E.; Huang, B. S.; Lowary, T. L.; Kimber, M. S.; Whitfield, C. Biosynthesis of a Conserved Glycolipid Anchor for Gram-Negative Bacterial Capsules. Nat. Chem. Biol. 2019, 15 (6), 632–640. 10.1038/s41589-019-0276-8.

(48) Jung, I. L.; Kim, I. G. Transcription of AhpC, KatG, and KatE Genes in Escherichia Coli Is Regulated by Polyamines: Polyamine-Deficient Mutant Sensitive to H2O2-Induced Oxidative Damage. Biochem. Biophys. Res. Commun. 2003, 301 (4), 915–922. 10.1016/S0006-291X(03)00064-0.

(49) Karamad, D.; Khosravi-Darani, K.; Khaneghah, A. M.; Miller, A. W. Probiotic Oxalate-Degrading Bacteria: New Insight of Environmental Variables and Expression of the Oxc and Frc Genes on Oxalate Degradation Activity. Foods. MDPI September 1, 2022. 10.3390/foods11182876.

(50) Liu, X.; Chang, Y.; Xu, Q.; Zhang, W.; Huang, Z.; Zhang, L.; Weng, S.; Leptihn, S.; Jiang, Y.; Yu, Y.; Hua, X. Mutation in the Two-Component Regulator BaeSR Mediates Cefiderocol Resistance and Enhances Virulence in Acinetobacter Baumannii. mSystems 2023, 8 (4). 10.1128/MSYSTEMS.01291-22.

(51) Prinz, W. A.; Åslund, F.; Holmgren, A.; Beckwith, J. The Role of the Thioredoxin and Glutaredoxin Pathways in Reducing Protein Disulfide Bonds in the Escherichia Coli Cytoplasm. Journal of Biological Chemistry 1997, 272 (25), 15661–15667. 10.1074/jbc.272.25.15661.

(52) Ryu, K. S.; Kim, C.; Kim, I.; Yoo, S.; Choi, B. S.; Park, C. NMR Application Probes a Novel and Ubiquitous Family of Enzymes That Alter Monosaccharide Configuration. Journal of Biological Chemistry 2004, 279 (24), 25544–25548. 10.1074/jbc.M402016200.

(53) Rhodius, V. A.; Suh, W. C.; Nonaka, G.; West, J.; Gross, C. A. Conserved and Variable Functions of the SigmaE Stress Response in Related Genomes. PLoS Biol. 2006, 4 (1), 0043–0059. 10.1371/JOURNAL.PBIO.0040002.

(54) Gibson, K. E.; Silhavy, T. J. The LysR Homolog LrhA Promotes RpoS Degradation by Modulating Activity of the Response Regulator SprE; 1999; Vol. 181. https://journals.asm.org/journal/jb.

(55) Ruelens, P.; de Visser, J. A. G. M. Clonal Interference and Mutation Bias in Small Bacterial Populations in Droplets. Genes (Basel). 2021, 12 (2), 1–10. 10.3390/genes12020223.

(56) Deatherage, D.E., Barrick, J.E.. (2014) Identification of mutations in laboratory-evolved microbes from next-generation sequencing data using breseq. Methods Mol. Biol. 1151:165–188.

