## Supplementary material for "Experimental evolution of collective β-lactam resistance in *Escherichia coli* via activation of a dormant outermembrane porin": SI_Ochoa_2026

**Table S1**. Wild-type strains genotype and sc-MIC for each of them.

| Strain | Genotype | sc-MIC (µg/mL) |
|---|---|---|
| TEM_Low | *REL606 galk::CFP pactem-G238S* | 1.0 |
| ΔompF_TEM_Low | *REL606 galk::CFP pactem-G238S, ΔompF* | 2.0 |
| TEM_High | *REL606 galk::CFP pactem- E104K M182T G238S T261M* | 96 |
| ΔompF_TEM_High | *REL606 galk::CFP pactem- E104K M182T G238S T261M, ΔompF* | 256 |

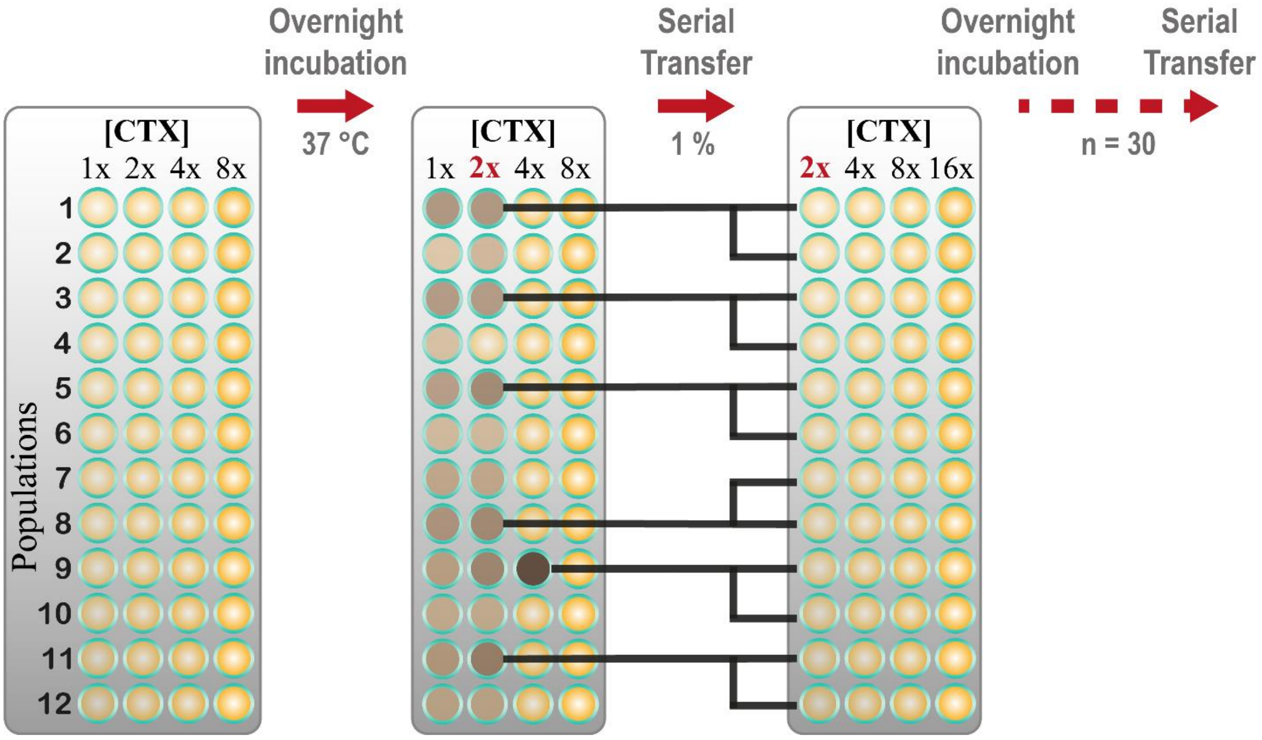


**Figure S1**. Selection strategy used for each transfer cycle during the experiment of evolution.

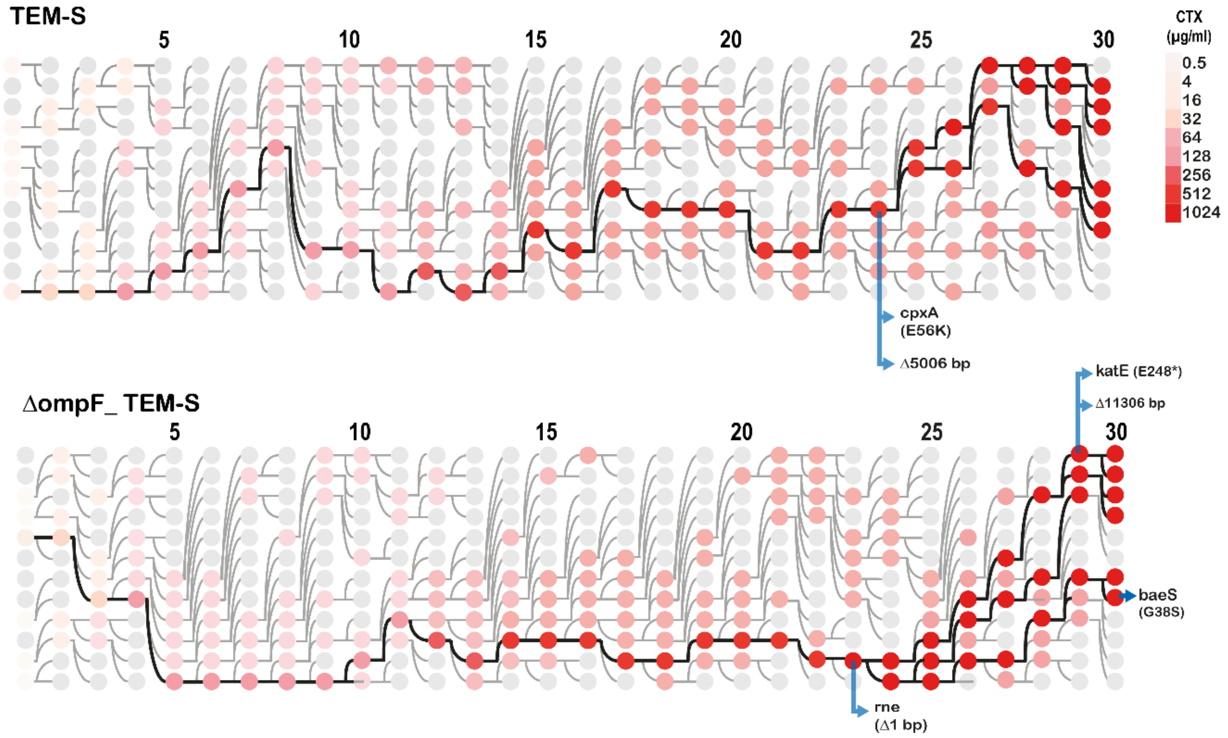


**Figure S2. Evolutionary dynamics and history of the six evolved lineages per strain** (see labels). Each column of circles represents the 12 lineages and their CTX tolerance (among the four concentrations applied each day); the colored circles indicate the lineages selected at each step, the highlighted colored circles and lines show the evolutionary history of the final six lineages selected for further analyses. Mutations identified in the six evolved final lineages of each ancestral strain are indicated using blue arrows. Mutations that are shared by two or more strains are indicated in the first common ancestor.

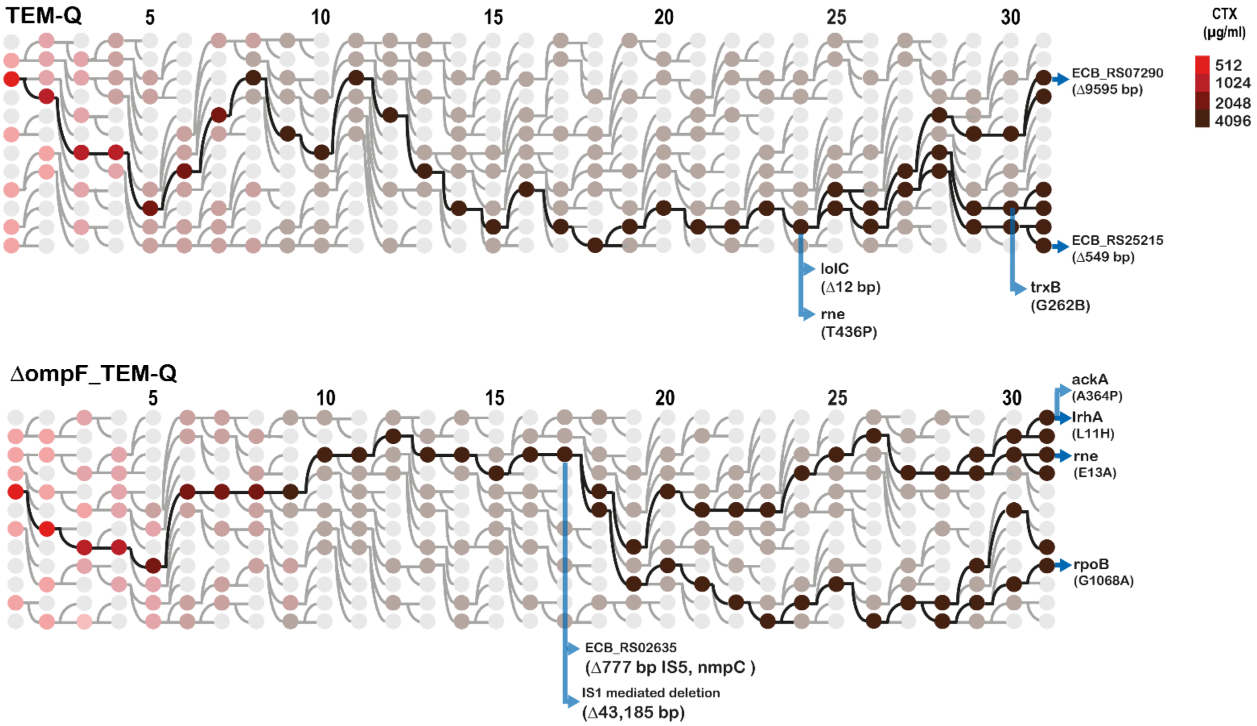


**Figure S3. Evolutionary dynamics and history of the six evolved lineages per strain** (see labels). Each column of circles represents the 12 lineages and their CTX tolerance (among the four concentrations applied each day); the colored circles indicate the lineages selected at each step, the highlighted colored circles and lines show the evolutionary history of the final six lineages selected for further analyses. Mutations identified in the six evolved final lineages of each ancestral strain are indicated using blue arrows. Mutations that are shared by two or more strains are indicated in the first common ancestor.

**Table S2**. Common deletions in evolved clones

| Strain | Range | Size (bp) | Event | Genes deleted |
|---|---|---|---|---|
| TEM_Low | 3,018,938-3,024,648 | 5,006 | IS1-mediated IS1A family | *[ECB_RS14895], ECB_RS14900, ECB_RS14905, ECB_RS14910, ECB_RS14915, ECB_RS14920, [ECB_RS14920]* |
| ΔompF_TEM_Low | 2,425,854-2,437,160 | 11,306 | IS-mediated | *[oxc], frc, yfdX, ypdI, yfdY, lpxP, ypdK, alaC, ypdA, ypdB, ypdC, [ptsP]* |
| TEM_High_P1 | 3,894,995-3,895,543 | 549 | IS3-like element IS150 family | *[ECB_RS25215], ECB_RS25215, [rbsD]* |
| TEM_High_P10 | 1,460,928-1,470,522 | 9,595 | IS1-like element IS1A family | *[ECB_RS07290], ECB_RS07290, cybB, ECB_RS07300, yncP, hokB, trg, ydcI, ydcJ, opgD, ydcH, rimL, [ydcK]* |
| ΔompF_TEM_High | 546,932-547,709 | 777 | IS1-like element IS1A family | *ECB_RS02630* |
| ΔompF_TEM_High | 1,460,928-1,504,112 | 43,185 | IS1-like element IS1A family | *[ECB_RS07290], ECB_RS07290, cybB, ECB_RS07300, yncP, hokB, trg, ydcI, ydcJ, opgD, ydcH, rimL, ydcK, tehA, tehB, ydcL, tnpA, insQ, ydcO, sutR, rlhA, yncJ, hicA, hicB, ydcR, ydcS, ydcT, ydcU, ydcV, patD, yncL, ortT, ydcY, ydcZ, mddA, curA, mcbR, pqqU, yncE, ansP, yncG, yncH, ECB_RS07505, [rhsD]* |

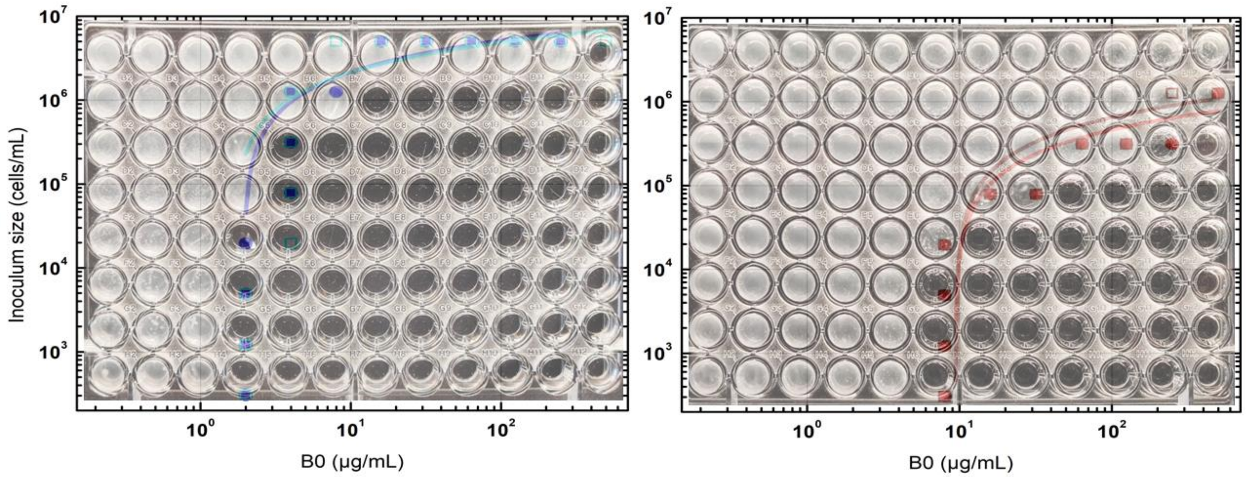


**Figure S4.** Typical result of the experiment assessing the dependence of MIC on the initial inoculum size for ancestral (left) and evolved (right) ΔompF_TEM-S populations. The X-axis shows the initial concentration of the antibiotic CTX (B0), and the Y-axis indicates the inoculum size. After 48 hours of incubation at 37 °C, bacterial proliferation is observed in wells with lower B0 values and is dependent on inoculum size. The MIC in each case is represented by a dot on the image. Two independent replicates were performed, each represented by a different color. Blue and red correspond to experiments with the ancestral and evolved strains, respectively. The solid lines represent the fit of the data to the model developed by Geyrhofer et al (reference 19 main manuscript).

**Table S3.** Statistical comparison of mean *T* values obtained from fitting the experimental data to Geyrhofer's mathematical model.

* Null Hypothesis: (Variance1/Variance2) = 1    Alternate Hypothesis: (Variance1/Variance2) ≠ 1

** Null Hypothesis: (mean1-mean2) = 0    Alternate Hypothesis: (mean1-mean2) ≠ 0

| | | | | | | | | | |
|---|---|---|---|---|---|---|---|---|---|
| | | **Tau (cells/mL)** | | | | | | | |
| | **n** | **TEM-Low_Anc** | **TEM-Low_Evol** | **ΔompF_TEM_Low_Anc** | **ΔompF_TEM-Low_Evol** | **TEM_High_Anc** | **TEM_High_Evol** | **ΔompF_TEM_High_Anc** | **ΔompF_TEM_High_Evol** |
| | 1 | 6.23 | 5.53 | 6.48 | 5.65 | 5.65 | 5.77 | 6.27 | 5.50 |
| | 2 | 6.42 | 6.22 | 6.43 | 5.80 | 5.69 | 6.17 | 5.90 | 5.07 |
| | 3 | 6.09 | 6.31 | 6.37 | 5.63 | 5.77 | 6.17 | 6.36 | 5.20 |
| | **mean** | **6.25E+00** | **6.01E+00** | **6.43E+00** | **5.69E+00** | **5.70E+00** | **6.03E+00** | **6.17E+00** | **5.26E+00** |
| | **SD** | **1.65E-01** | **4.26E-01** | **5.28E-02** | **9.26E-02** | **5.74E-02** | **2.31E-01** | **2.43E-01** | **2.21E-01** |
| *Levene's test | F-value | 4.54 | equal variance | 1.96 | equal variance | 8.43 | different variance | 0.08 | equal variance |
| | p-value | 0.1 | | 0.23 | | 0.044 | | 0.79 | |
| **T-test | t-value | 0.86 | mean values are NOT different | 11.93 | mean values ARE different | -2.41 | mean values are NOT different | 4.8 | mean values ARE different |
| | p-value | 0.44 | | 0.0003 | | 0.12 | | 0.0084 | |

**Table S4**. Statistical comparison of mean *T* values obtained from fitting the experimental data to Geyrhofer's mathematical model.

| | | | | | | | | | |
|---|---|---|---|---|---|---|---|---|---|
| | | **μ (μg/mL)** | | | | | | | |
| | **n** | **TEM-Low_Anc** | **TEM-Low_Evol** | **ΔompF_TEM_Low_Anc** | **ΔompF_TEM-Low_Evol** | **TEM_High_Anc** | **TEM_High_Evol** | **ΔompF_TEM_High_Anc** | **ΔompF_TEM_High_Evol** |
| | 1 | 0.12 | 0.79 | 0.29 | 0.97 | 1.88 | 2.24 | 2.20 | 1.98 |
| | 2 | -0.08 | 1.26 | 0.22 | 0.97 | 1.97 | 2.28 | 2.49 | 1.78 |
| | 3 | 0.13 | 1.05 | 0.35 | 0.99 | 2.15 | 2.28 | 2.41 | 1.92 |
| | **mean** | **5.90E-02** | **1.01E+00** | **2.78E-01** | **9.79E-01** | **2.00E+00** | **2.27E+00** | **2.36E+00** | **1.89E+00** |
| | **SD** | **1.19E-01** | **2.39E-01** | **6.40E-02** | **1.18E-02** | **1.38E-01** | **2.60E-02** | **1.53E-01** | **1.02E-01** |
| *Levene's test | F-value | 0.92 | equal variance | 2.88 | equal variance | 4.79 | equal variance | 0.73 | equal variance |
| | p-value | 0.39 | | 0.17 | | 0.09 | | 0.44 | |
| **T-test | t-value | -6.32 | mean values ARE different | -18.53 | mean values ARE different | -3.27 | mean values ARE NOT different | 4.45 | mean values ARE different |
| | p-value | 0.0032 | | 4.99E-05 | | 0.075 | | 0.01 | |

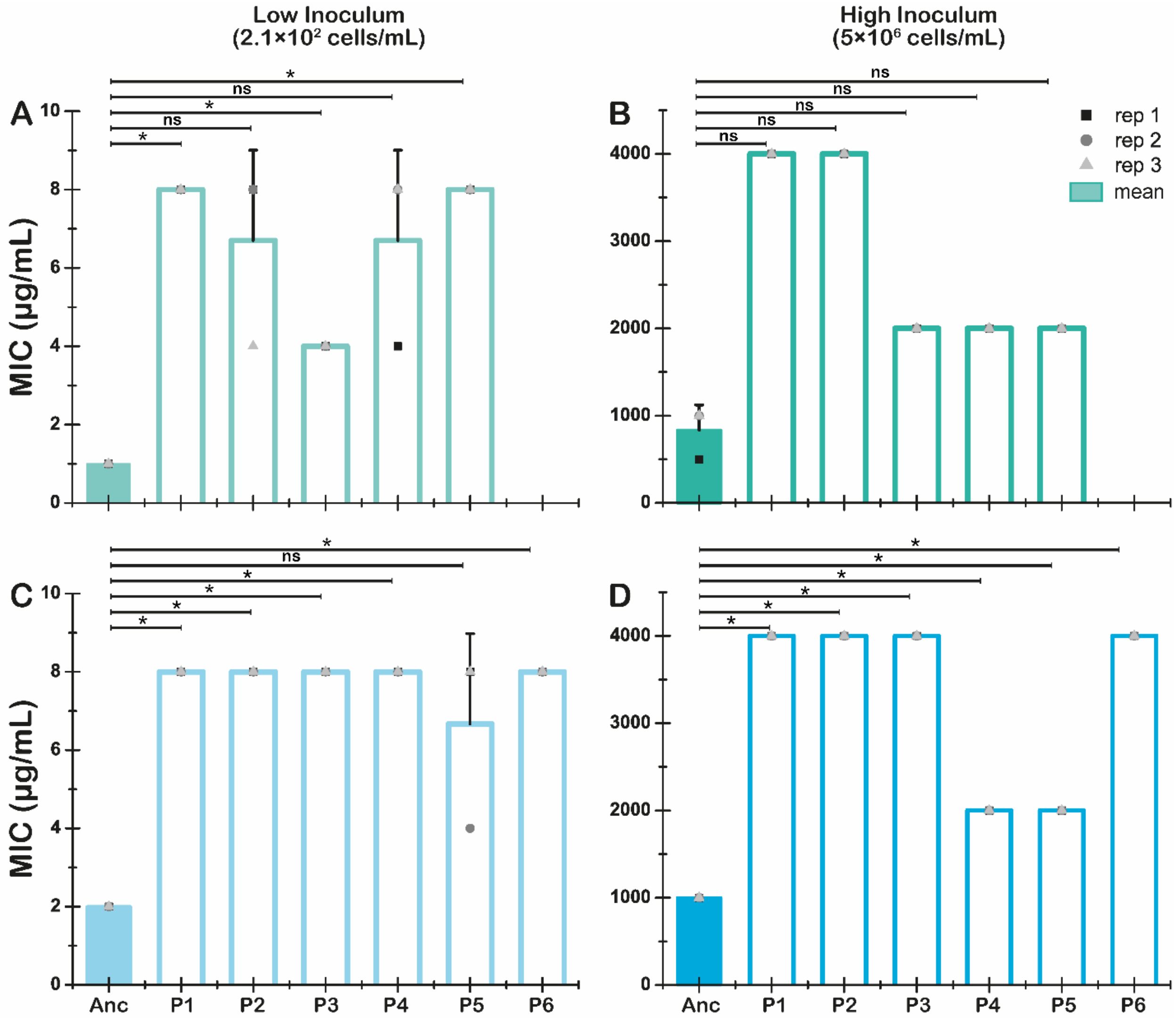


**Figure S5**. MIC evaluated at high an low inculum sizes for TEM-S and ΔompF_TEM-S wild-type and evolved lineages.

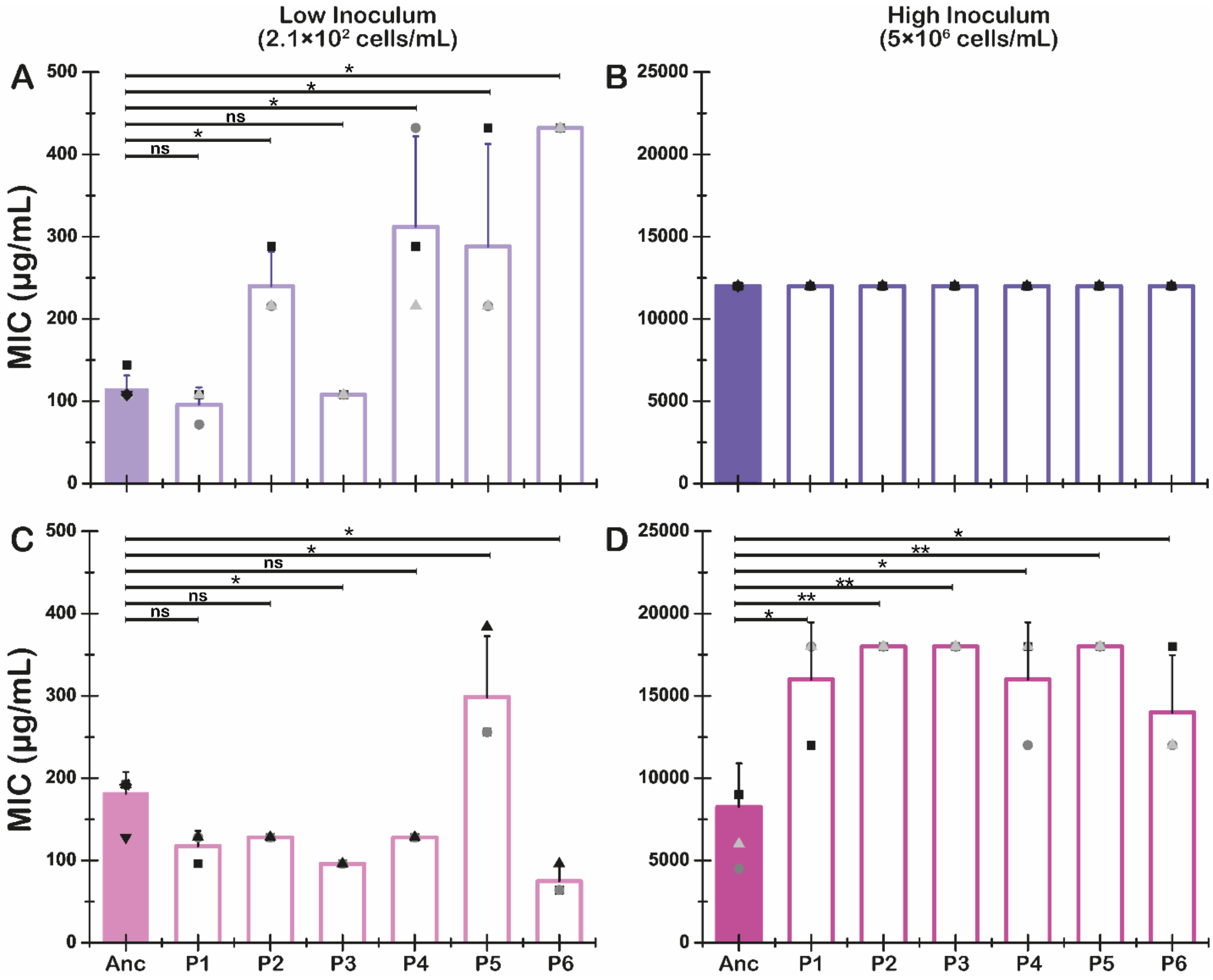


**Figure S6**. MIC evaluated at high an low inculum sizes for TEM-Q and ΔompF_TEM-Q wild-type and evolved lineages.

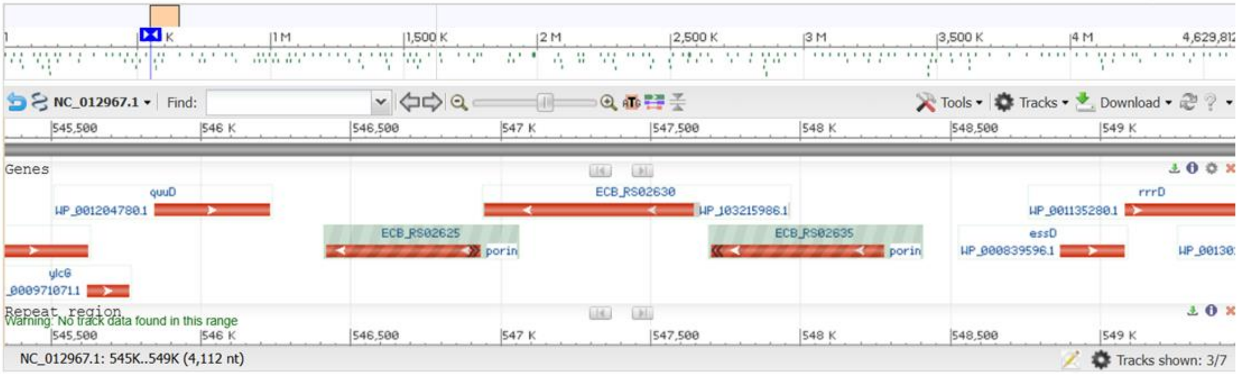


NC_012967.1: Escherichia coli B str. REL606, complete sequence (November 21, 2025)

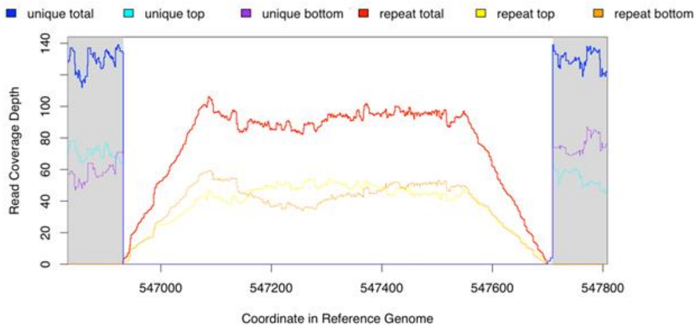


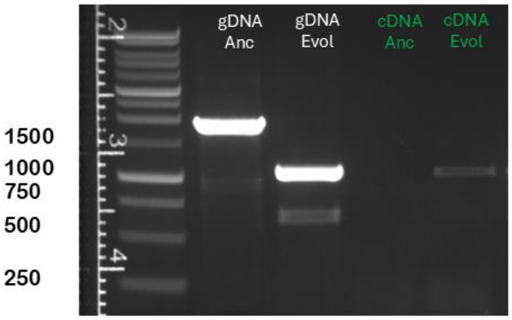


Deatherage, D.E., Barrick, J.E. (2014), breseq. Methods Mol. Biol. 1151: 165–188

**Figure S7**. Porin activation during the evolution experiment via the deletion of an IS1A (*ECB_RS0263)* element in between two fragments that together codified for a porin annotated as *nmpC* in the reference genome of *E. coli* line used in this study (REL606, NC_012967.1).

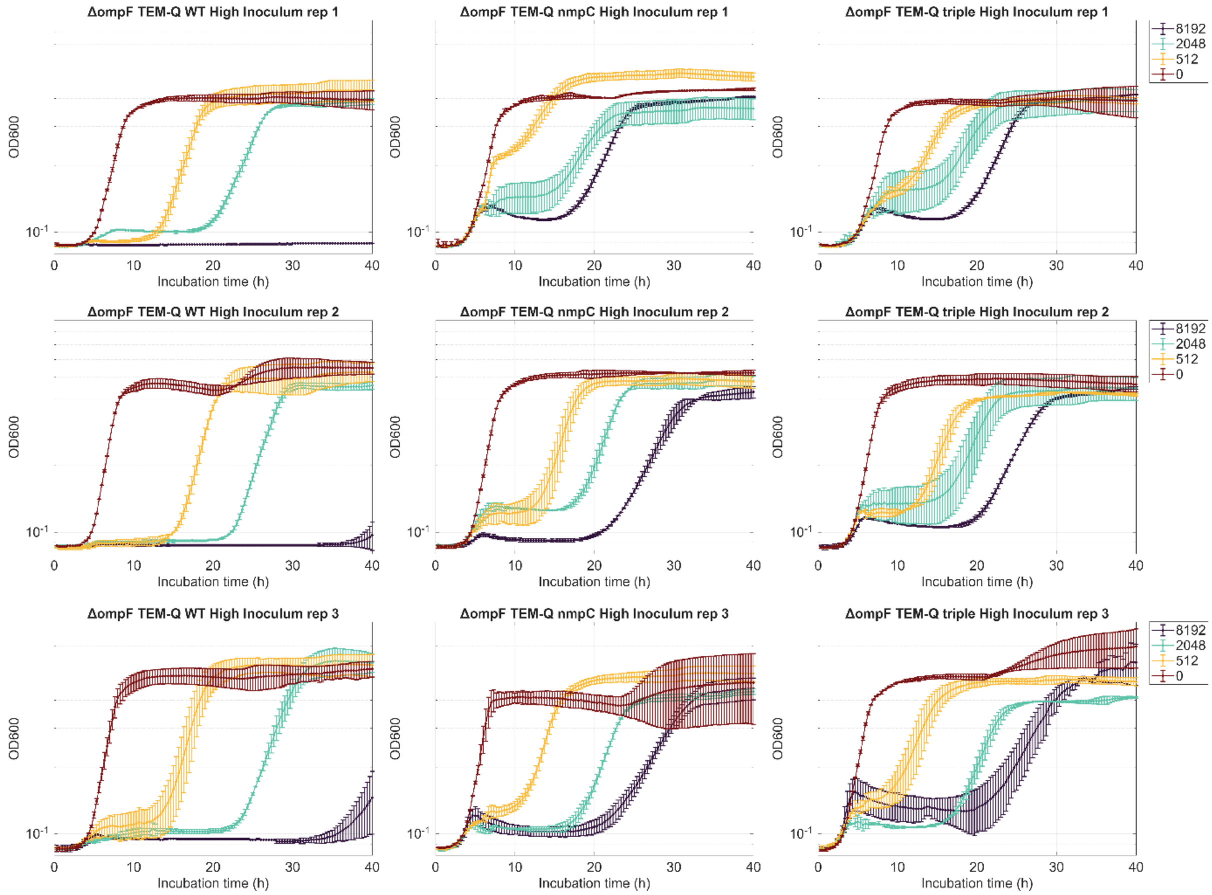


**Figure S8.** Growth curve of ΔompF_TEM-Q evolutionary landscape evaluated at high inoculum ($5\times10^6$ cells/mL). Results are show for three different biological replicates (rows) and for each member of the landscape (columns). The error bars in each plots correspond to the standard deviation of two technical replicates for each biological replicate.

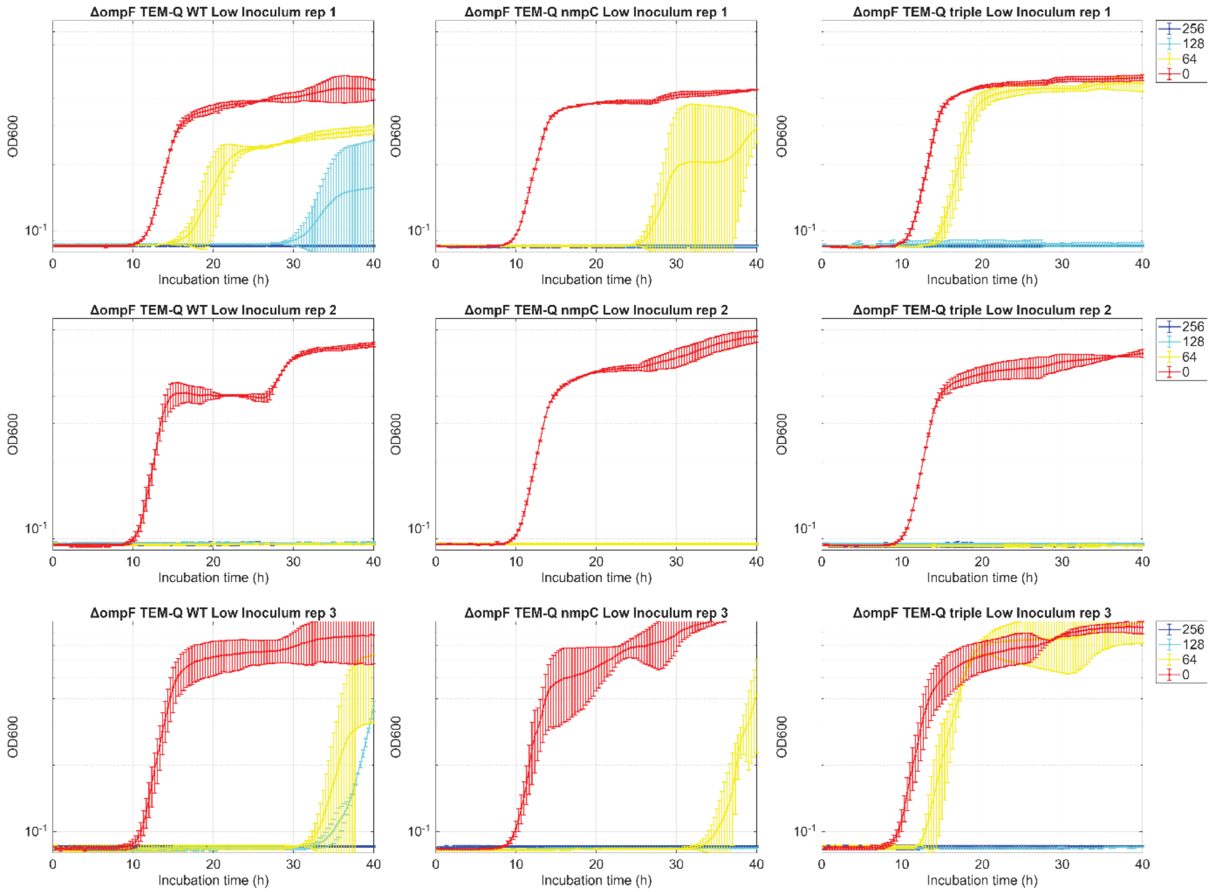


**Figure S9.** Growth curve of ΔompF_TEM-Q evolutionary landscape evaluated low inoculum size ($5 \times 10^2$ cells/mL). Results are show for three different biological replicates (rows) and for each member of the landscape (columns). The error bars in each plots correspond to the standard deviation of two technical replicates for each biological replicate.

**Table S5**. Mann–Whitney test results comparing the kinetic parameters of ΔompF-TEM-Q (wt) with its derived evolved lineages, as well as comparisons among the evolved lineages. N is equal to 6 replicates, and significance was evaluated at α = 0.05. Boxes shaded in green indicate significant difference, whereas red indicates no significant difference. Results are shown for bacterial cultures initiated at 5 × $10^6$ cells/mL. NA means Not Applicable.

| Bacteria culture started at High Inoculum (5×$10^6$ cells/mL) | | | | |
|---|---|---|---|---|
| Initial [CTX] | Compared strains | Filamentation rate | recovery rate | recovery time |
| | | p value (α=0.05) | p value (α=0.05) | p value (α=0.05) |
| 8192 | wt-nmpc | 0.025 | 0.019 | 0.024 |
| 4096 | wt-nmpc | 0.008 | 0.066 | 0.044 |
| 2048 | wt-nmpc | 0.005 | 0.3 | 0.005 |
| 512 | wt-nmpc | 0.005 | 0.065 | 0.008 |
| 0 | wt-nmpc | NA | 1 | 0.19 |
| 8192 | wt-triple | 0.005 | 0.012 | 0.023 |
| 4096 | wt-triple | 0.009 | 0.066 | 0.013 |
| 2048 | wt-triple | 0.005 | 0.031 | 0.005 |
| 512 | wt-triple | 0.005 | 0.005 | 0.008 |
| 0 | wt-triple | NA | 0.81 | 0.292 |
| 8192 | nmpC-triple | 0.936 | 1 | 0.808 |
| 4096 | nmpC-triple | 0.091 | 0.422 | 0.335 |
| 2048 | nmpC-triple | 0.378 | 0.378 | 0.637 |
| 512 | nmpC-triple | 0.469 | 0.378 | 0.936 |
| 0 | nmpC-triple | NA | 0.575 | 0.935 |

**Table S6**. Mann–Whitney test results comparing the kinetic parameters of ΔompF-TEM-Q (wt) with its derived evolved lineages, as well as comparisons among the evolved lineages. N is equal to 6 replicates, and significance was evaluated at α = 0.05. Boxes shaded in green indicate significant difference, whereas red indicates no significant difference. Results are shown for bacterial cultures initiated at 5 × $10^2$ cells/mL.

| Bacteria culture started at Low Inoculum (5×$10^2$ cells/mL) | | | |
|---|---|---|---|
| Initial [CTX] | Compared strains | Filamentation rate | recovery rate |
| | | p value (α=0.05) | p value (α=0.05) |
| 64 | wt-nmpc | 0.194 | 0.112 |
| 0 | wt-nmpc | 0.936 | 0.0122 |
| 64 | wt-triple | 0.112 | 0.052 |
| 0 | wt-triple | 0.936 | 0.372 |
| 64 | nmpC-triple | 0.471 | 0.03 |
| 0 | nmpC-triple | 0.422 | 0.196 |

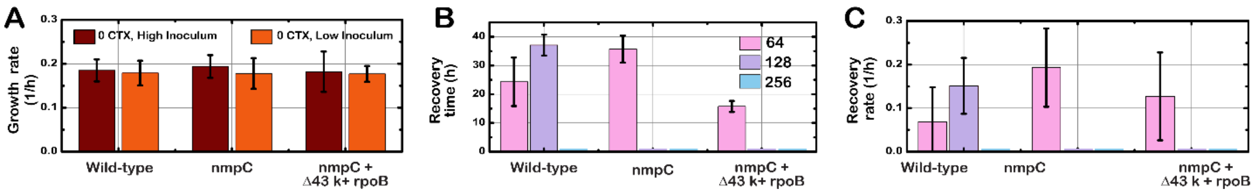


**Figure S10**. Parameters associated to the growth curves for the three selected members of the evolutionary landscape of ΔompF_TEM-Q. (A) Growth rate in the absence of antibiotic; (B) recovery time at low inoculum (LI); (C) recovery rate at LI.

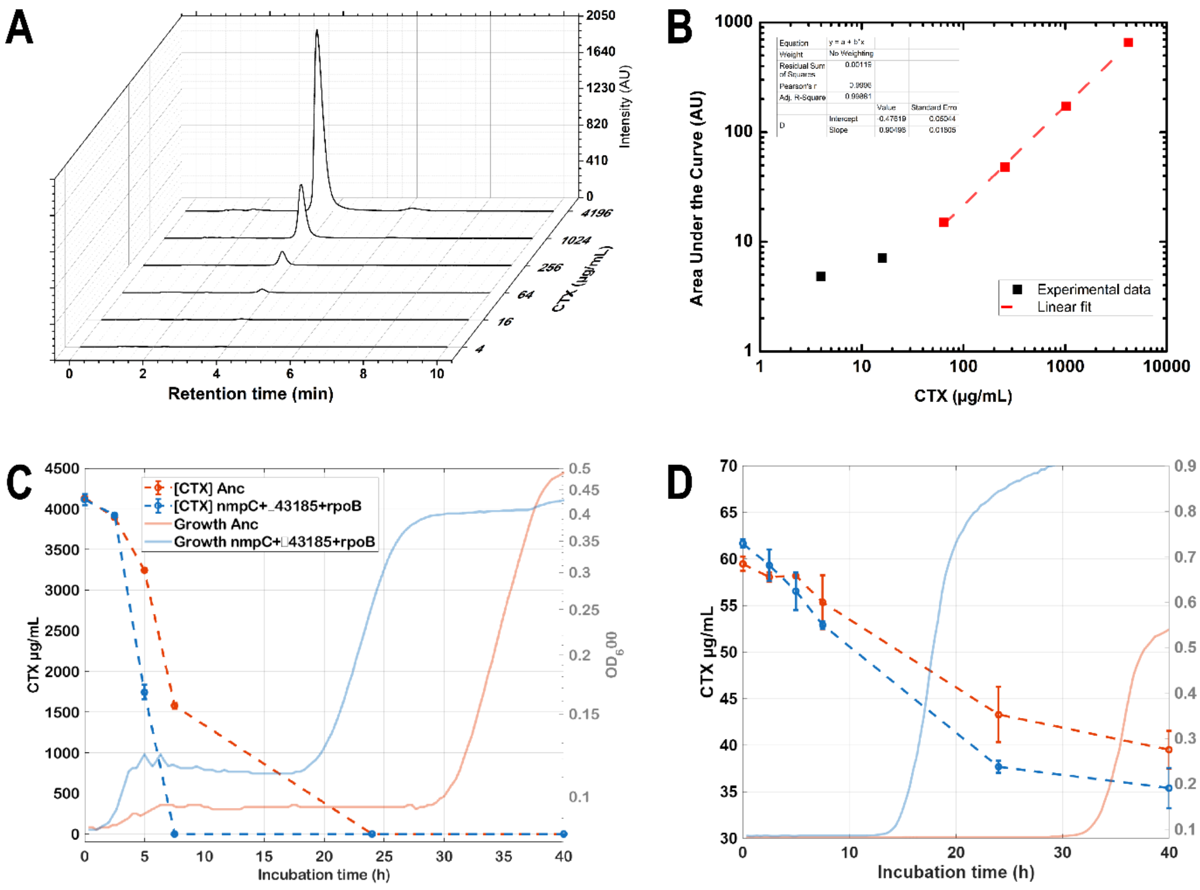

**Figure S11.** (A) Representative chromatograms of a standard solution of Cefotaxime injected into the HPLC system to build a calibration curve. (B) Area under the curve (AUC) corresponding to the CTX peak in the HPLC system as a function of CTX concentration. (C) Degradation rate of CTX over time in bacterial cultures of the ΔompF_TEM_High and its evolved lineage, the triple mutant (nmpC + Δ43,184 bp + rpoB). Cultures were initiated at $5\times10^6$ cells/mL and 4096 μg/mL of CTX. (D) Cultures were started at $5\times10^2$ cells/mL and 64 μg/mL of CTX. The left axis shows CTX concentration over incubation time. Each data point represents the mean of two technical replicates, and error bars indicate the standard deviation. The right axis shows the growth curves recorded for the bacterial cultures from which samples were taken for HPLC analysis.
